# (*R,S*)-ketamine’s rapid-acting antidepressant effects are modulated by NR2B-containing NMDA receptors on adult-born hippocampal neurons

**DOI:** 10.1101/2023.11.28.569043

**Authors:** Nicholas E. Bulthuis, Josephine C. McGowan, Liliana R. Ladner, Christina T. LaGamma, Sean C. Lim, Claire X. Shubeck, Rebecca A. Brachman, Ezra Sydnor, Ina P. Pavlova, Dong-oh Seo, Michael R. Drew, Christine A. Denny

## Abstract

Standard antidepressant treatments often take weeks to reach efficacy and are ineffective for many patients. (*R,S*)-ketamine, an *N*-methyl-D-aspartate (NMDA) antagonist, has been shown to be a rapid-acting antidepressant and to decrease depressive symptoms within hours of administration. While previous studies have shown the importance of the NR2B subunit of the NMDA receptor (NMDAR) on interneurons in the medial prefrontal cortex (mPFC), no study has investigated the influence of NR2B-expressing adult-born granule cells (abGCs). In this study, we examined whether (*R,S*)-ketamine’s efficacy depends upon these adult-born hippocampal neurons using a genetic strategy to selectively ablate the NR2B subunit of the NMDAR from Nestin^+^ cells. To validate our findings, we also used several other transgenic lines including one in which NR2B was deleted from an interneuron (Parvalbumin (PV)^+^) population. We report that in male mice, NR2B expression on 6-week-old adult-born neurons is necessary for (*R,S*)-ketamine’s effects on behavioral despair in the forced swim test (FST) and on hyponeophagia in the novelty suppressed feeding (NSF) paradigm, as well on fear behavior following contextual fear conditioning (CFC). In female mice, NR2B expression is necessary for effects on hyponeophagia in the NSF. We also find that ablating neurogenesis increases fear expression in CFC, which is buffered by (*R,S*)-ketamine administration. In line with previous studies, these results suggest that 6-week-old adult-born hippocampal neurons expressing NR2B partially modulate (*R,S*)-ketamine’s rapid-acting effects. Future work targeting these 6-week-old adult-born neurons may prove beneficial for increasing the efficacy of (*R*,*S*)-ketamine’s antidepressant actions.

## INTRODUCTION

Major depressive disorder (MDD) is a debilitating psychiatric disorder that affects approximately 17% of the U.S. population [1]. The World Health Organization (WHO) has declared MDD to be the leading cause of disability worldwide and a major contributor to the global burden of disease [2] far earlier than the previously projected year of 2030 [3]. Yet, the cellular and molecular mechanisms underlying the disorder have not yet been fully revealed, and current antidepressant medications often take weeks to take effect and may not be effective in some patients [4]. Especially for individuals who are at risk for suicide, rapid-acting and long-lasting treatments for MDD are long overdue.

(*R,S*)-ketamine, an *N*-methyl-D-aspartate receptor (NMDAR) antagonist, has been shown to be a rapid-acting antidepressant, showing effects within 2 hours after a single dose that can last up to 3 weeks [5] even in individuals with treatment-resistant depression (TRD). Additionally, we and others have found that (*R,S*)-ketamine promotes resilience as a prophylactic against stress-induced behaviors, such as behavioral despair, learned fear, and hyponeophagia [6–12]. However, to develop therapeutics that lack the dissociative and addictive properties of (*R,S*)-ketamine [13], and to further clarify the biological underpinnings of MDD, we must better understand how (*R,S*)-ketamine confers its antidepressant and prophylactic effects.

Previous research suggests that the glutamatergic system, and specifically NMDARs, are important to the mechanism of action of antidepressant treatments [14, 15]. The NMDAR is a heteromultimeric complex that consists of two obligatory NR1 subunits and two NR2 subunits, which are encoded by four genes (NR2A-D) [16]. NR2B has been identified to form the glutamate-binding pocket of NMDARs, indicating that (*R,S*)-ketamine may interact directly with NR2B to modulate channel gating [17]. A reduction in human NR2B subunit expression was found in the prefrontal cortex (PFC) of patients with MDD as compared to controls [18], and 3 potentially functional SNPs that are considered risk variables for MDD and predictors of TRD were identified in the NR2B gene *GRIN2B* [19]. Furthermore, NR2B on principal cortical neurons is necessary for (*R,S*)-ketamine’s effects in mice [20]. Collectively, these data suggest that NR2B is a critical mediator of antidepressant efficacy. However, while drugs selective for NR2B have shown efficacy in both preclinical and clinical studies [21–23], these results are not consistent [24, 25].

Adult hippocampal neurogenesis, or the process by which neurons are generated from neural stem cells in adulthood, has also been implicated in antidepressant treatment. MDD patients treated with antidepressants have increased neural progenitor density in the anterior dentate gyrus (DG) of the hippocampus when compared to MDD patients without antidepressant treatment [26]. While this study and others [27–30] suggest a critical role for neurogenesis in MDD, only antidepressants currently on the market were examined. While (*R,S*)-ketamine has previously been found to increase neurogenesis at repeated doses [31], and the activity of adult-born granule cells (abGCs) is both necessary and sufficient for its rapid-acting antidepressant effects [32], no studies have specifically probed the role of NR2B on abGCs in mediating (*R,S*)-ketamine’s antidepressant efficacy.

Here, we investigated how the NR2B subunit in concert with adult hippocampal neurogenesis modulates (*R,S*)-ketamine’s effectiveness. We hypothesized that NR2B-containing abGCs at 6 weeks of age are important for the effects of (*R,S*)-ketamine, as we and others have previously shown that these neurons exhibit heightened synaptic plasticity [33] and influence both mood and cognition [34–38]. Here, we generated a triple transgenic mouse line, NestinCreER^T2^ x NR2B flox/flox (NR2B^f/f^) x R26R-STOP-floxed-enhanced yellow-fluorescent protein (EYFP), which allows for the tamoxifen (TAM)-inducible deletion of NR2B from Nestin^+^ abGCs. We found that deletion of NR2B from Nestin^+^ neurons at 6 but not 2 weeks of age occludes the antidepressant effects of (*R,S*)-ketamine. To validate these findings, we used a separate genetic line to arrest adult hippocampal neurogenesis, which resulted in increased fear expression that was buffered by (*R,S*)-ketamine administration. Overall, our results suggest that NR2B-expressing 6-week-old abGCs partially modulate (*R,S*)-ketamine’s rapid-acting effects. These data propose a potential mechanism of (*R,S*)-ketamine action and reveal a novel target for antidepressant development.

## MATERIALS AND METHODS

### Mice

Mice were housed 4–5 per cage in a 12-h (06:00-18:00) light-dark colony room at 22°C. Food and water were provided *ad libitum*. Behavioral testing was performed during the light phase. All experiments were approved by the Institutional Animal Care and Use Committees (IACUCs) at Columbia University Irving Medical Center (CUIMC) and the New York State Psychiatric Institute (NYSPI). All other mouse genotyping information can be found in the **Supplement**.

### Drugs

*Tamoxifen (TAM):* NestinCreER^T2^ mice (8- to 10-weeks-old) were injected with 3 mg of TAM in a vehicle (Veh) solution of corn oil/10% EtOH or a Veh solution intraperitoneally (i.p.) once per day for 5 consecutive days.

*(R,S)-ketamine:* A single injection of saline (0.9% NaCl) or (*R,S*)-ketamine (10 or 30 mg/kg) (Ketaset III, Ketamine HCl injection, Fort Dodge Animal Health, Fort Dodge, IA) was administered 2 hours before the start of the forced swim test (FST) for all groups. The dose was based on prior studies showing sex-specific dosing effects of (*R,S*)-ketamine [6–12]. (*R,S*)-ketamine was prepared in physiological saline and all injections were administered i.p. in volumes of 0.1 cc per 10 mg body weight.

### Statistical Analysis

All data were analyzed using Prism v9.0.0 (GraphPad, San Diego, CA) with alpha set to 0.05. Results from analyses are expressed as means ± SEM. * p < 0.05, ** p < 0.01, *** p < 0.001, **** p < 0.0001. Effects of Drug, Genotype, Treatment, and Time were analyzed using an analysis of variance (ANOVA) or repeated measures ANOVA (RMANOVA) with a Greenhouse-Geisser correction where appropriate. Significant ANOVAs were followed with a Šídák’s post-hoc analysis or unpaired *t*-tests as indicated. For the novelty suppressed feeding (NSF) paradigm, statistical differences were determined using a Kaplan-Meier survival analysis. All statistical tests and p-values are listed throughout the text and in **Supplemental Table 1**.

**All other Methods are listed in the Supplement.**

## RESULTS

### Deletion of NR2B from abGCs less than 6 weeks old occludes (*R,S*)-ketamine’s effects on hyponeophagia and fear expression in male mice

To determine if the NR2B subunit is required for (*R,S*)-ketamine’s effects, we generated a triple transgenic mouse line that enabled EYFP labeling and deletion of NR2B in Nestin^+^ abGCs (**Fig. 1a**). Here, male mice were administered 5 daily injections of Veh or TAM. Six weeks later, mice were administered a single injection of saline or (*R,S*)-ketamine (30 mg/kg) (**Fig. 1b**). Two hours later, mice were administered day 1 of the forced swim test (FST) followed by several other assays to measure behavioral despair, locomotion, hyponeophagia, and fear behavior. Robust EYFP labeling in the dentate gyrus (DG) of the hippocampus (HPC) indicated genetic deletion of NR2B (**Fig. 1c**).

**Fig. 1.**
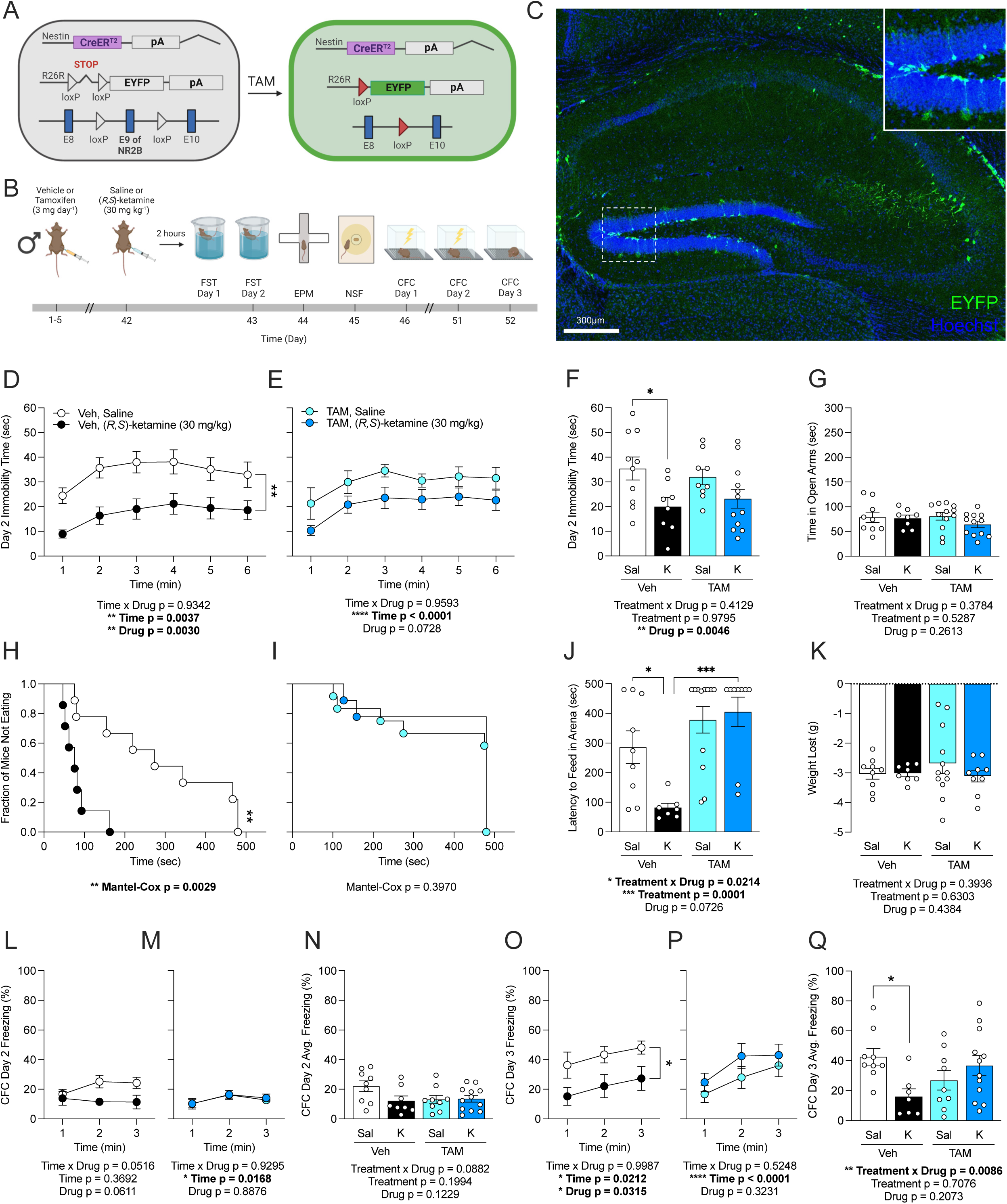
Deleting NR2B from 6-week-old adult-born hippocampal neurons occludes (*R,S*)-ketamine’s antidepressant effects in male mice. (**A**) Genetic schematic. (**B**) Experimental timeline. (**C**) Representative section showing EYFP (green) and Hoechst (blue) labeling in the HPC of NestinCreER^T2^ x NR2B x EYFP male mice. (**D**) Veh-K mice exhibited reduced immobility time in the FST as compared to Veh-Sal mice. (**E**) TAM-Sal and TAM-K mice exhibited comparable immobility time. (**F**) Average immobility time in the FST was decreased in Veh-K mice as compared to Veh-Sal mice, while K had no effect in TAM mice. (**G**) Time spent in the open arms of the EPM did not differ across groups. (**H**) In the NSF, Veh-K mice had a decreased latency to approach the pellet when compared to Veh-Sal mice. (**I**) TAM-Sal and TAM-K mice exhibited comparable latencies. (**J**) The latency to feed in the NSF was significantly reduced in Veh-K as compared to Veh-Sal mice. This effect was ablated in TAM mice. (**K**) Weight loss in the NSF did not differ across groups. (**L-N**) On day 2 of CFC, the percentage of freezing did not differ between the groups. (**O**) On day 3 of CFC, Veh-K mice froze significantly less than Veh-Sal mice. (**P**) There was no difference in freezing between TAM-Sal or TAM-K mice. (**Q**) Average freezing during CFC Day 3 was decreased in Veh-K mice as compared to Veh-Sal mice, while K had no effect in TAM mice. (n = 8-12 male mice / group). Error bars represent ± SEM. * p < 0.05; ** p < 0.01; *** p < 0.001; **** p < 0.0001. HPC, hippocampus; EYFP, enhanced yellow fluorescent protein; Veh, vehicle; TAM, tamoxifen; FST, forced swim test; EPM, elevated plus maze; NSF, novelty suppressed feeding; CFC, contextual fear conditioning; sec, seconds; min, minutes; h, hours; mg, milligrams; kg, kilograms; g, grams; Sal, saline; K, (*R,S*)-ketamine (30 mg/kg).

On day 1 of the FST, neither Drug nor Treatment impacted immobility time (**Fig. S1a-1c**). On day 2 of the FST, Veh-treated mice administered (*R,S*)-ketamine spent less time immobile than saline-injected Veh-treated mice (**Fig. 1d**). However, this difference was abolished between saline- and (*R,S*)-ketamine-injected TAM-treated mice (**Fig. 1e**). There was an effect of Drug but not of Treatment or an interaction on the average immobility time (**Fig. 1f**). These data indicate that deletion of the NR2B subunit on 6-week-old abGCs occludes the (*R,S*)-ketamine effect on behavioral despair.

Next, we assayed mice in the elevated plus maze (EPM). There was no effect of Drug, Treatment, or an interaction on time spent in the open (**Fig. 1g**) or closed arms (**Fig. S1d**). In the novelty suppressed feeding (NSF) paradigm, (*R,S*)-ketamine-injected Veh-treated mice displayed decreased latency to eat as compared with saline-injected Veh-treated mice (**Fig. 1h**). However, TAM treatment abolished this difference between saline- and (*R,S*)-ketamine-injected mice (**Fig. 1i-1j**). There were no differences in weight loss (**Fig. 1k**). These data indicate that deletion of the NR2B subunit on 6-week-old abGCs occludes (*R,S*)-ketamine’s effects on hyponeophagia.

Mice were then administered a 2-trial contextual fear conditioning (CFC) paradigm. On day 1 (**Fig. S1e-1g**) and day 2 (**Fig. 1l-1n**) of CFC, (*R,S*)-ketamine-injected Veh-treated mice trended towards freezing less than saline-injected Veh-treated mice (p = 0.0516). However, on day 3 of CFC, (*R,S*)-ketamine-injected Veh-treated mice displayed less freezing than saline-injected Veh-treated mice (**Fig. 1o**). Notably, this difference was abolished between saline- and (*R,S*)-ketamine-injected TAM-treated mice (**Fig. 1p-1q**). These data suggest that deletion of NR2B on 6-week-old abGCs occludes (*R,S*)-ketamine’s effects in contextual fear expression.

### Deletion of NR2B from 6-week-old abGCs selectively occludes (*R,S*)-ketamine’s effects on hyponeophagia in female mice

To extend our findings from male mice, we next deleted NR2B in abGCs as described in **Fig. 1**, but in female NestinCreER^T2^ x NR2B^f/f^ x EYFP^f/f^ mice (**Fig. 2a-2c**). For these and all subsequent female data, (*R,S*)-ketamine was administered at 10 mg/kg, which we have previously shown to be the effective dose in females [11, 39]. On day 1 (**Fig. S2a-2c**) and day 2 of the FST (**Fig. S2d-2f**), neither Drug nor Treatment impacted immobility time. These data indicate that unlike in male mice, (*R,S*)-ketamine does not produce robust effects on behavioral despair in females.

**Fig 2.**
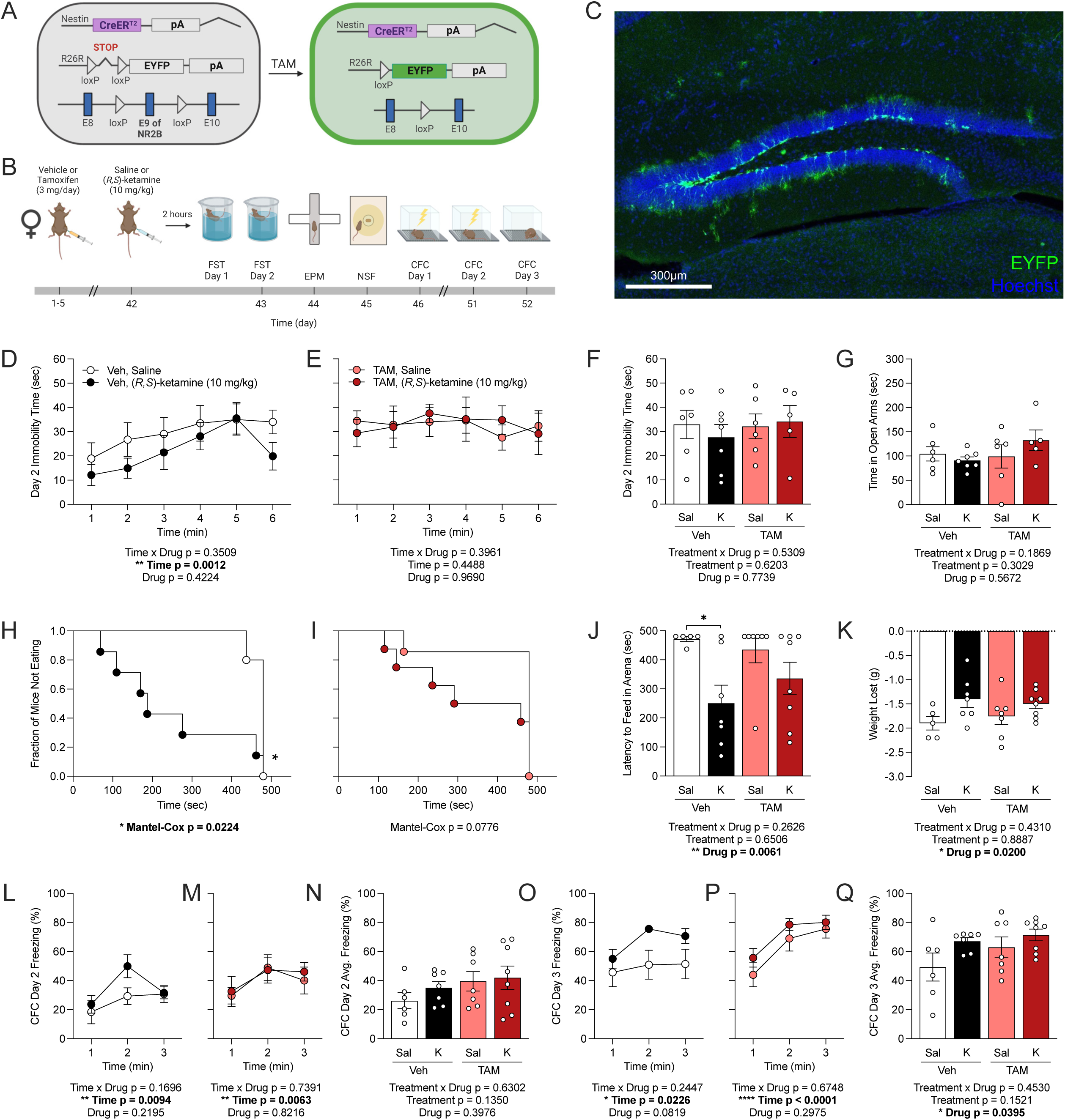
Deletion of NR2B from 6-week-old adult-born neurons selectively occludes (*R,S*)-ketamine’s effects in female mice. (**A**) Genetic schematic. (**B**) Experimental timeline. (**C**) Representative section showing EYFP (green) and Hoechst (blue) labeling in the HPC of NestinCreER^T2^ x NR2B^f/f^ x EYFP female mice. (**D-F**) There was no difference in immobility time between all groups on day 2 of the FST. (**G**) Time spent in the open arms of the EPM did not differ across groups. (**H**) In the NSF, Veh-K mice had a decreased latency to approach the pellet when compared to Veh-Sal mice. (**I**) TAM-Sal and TAM-K mice exhibited comparable latencies. (**J**) The latency to feed in the NSF was significantly reduced in Veh-K as compared to Veh-Sal mice. This effect was ablated in TAM mice. (**K**) There was a main effect of Drug on weight lost. (**L-N**) On day 2 of CFC, all groups froze at a comparable level. (O-Q) On day 3 of CFC, there was a main effect of Drug on freezing. (n = 5-8 female mice / group). Error bars represent ± SEM. * p < 0.05; ** p < 0.01; **** p < 0.0001. HPC, hippocampus; EYFP, enhanced yellow fluorescent protein; Veh, vehicle; TAM, tamoxifen; FST, forced swim test; EPM, elevated plus maze; NSF, novelty suppressed feeding; CFC, contextual fear conditioning; sec, seconds; min, minutes; h, hours; mg, milligrams; kg, kilograms; g, grams; Sal, saline; K, (*R,S*)-ketamine (10 mg/kg).

In the EPM, there was no effect of Drug, Treatment, or an interaction on time in the open (**Fig. 2g**) or closed arms (**Fig. S2d**). In the NSF, (*R,S*)-ketamine-injected Veh-treated mice displayed decreased latency to feed as compared with saline-injected Veh-treated mice (**Fig. 2h**). However, this difference was abolished in TAM-treated mice (**Fig. 2i-2j**). There was an overall effect of Drug on weight loss (**Fig. 2k**). These data indicate that deletion of NR2B on 6-week-old abGCs in female mice occludes (*R,S*)-ketamine’s effects on hyponeophagia.

On day 1 (**Fig. S2e-2g**) and day 2 of CFC (**Fig. 2l-2n**), there was no effect of Drug or Treatment on freezing behavior. On day 3 of CFC (**Fig. 2o-2q**), there an effect of Drug on average freezing behavior, but no post-hoc tests reached significance, indicating no effect of (*R,S*)-ketamine on fear behavior in females. These data suggest that in female mice, deletion of NR2B selectively occludes (*R,S*)-ketamine’s effects on hyponeophagia.

### Deletion of NR2B from abGCs less than 2 weeks old selectively influences (*R,S*)-ketamine’s fear effects in male mice

To validate that the occlusion of (*R,S*)-ketamine’s effects via NR2B is specific to 6-week-old abGCs, we next deleted NR2B from 2-week-old neurons in NestinCreER^T2^ x NR2B^f/f^ x EYFP^f/f^ male mice (**Fig. 3a**). Mice were administered 5 injections of Veh or TAM and then a single injection of saline or (*R,S*)-ketamine (30 mg/kg) 2 weeks later (**Fig. 3b**). Robust EYFP labeling in the DG indicated genetic deletion of NR2B (**Fig. 2c**).

**Fig. 3.**
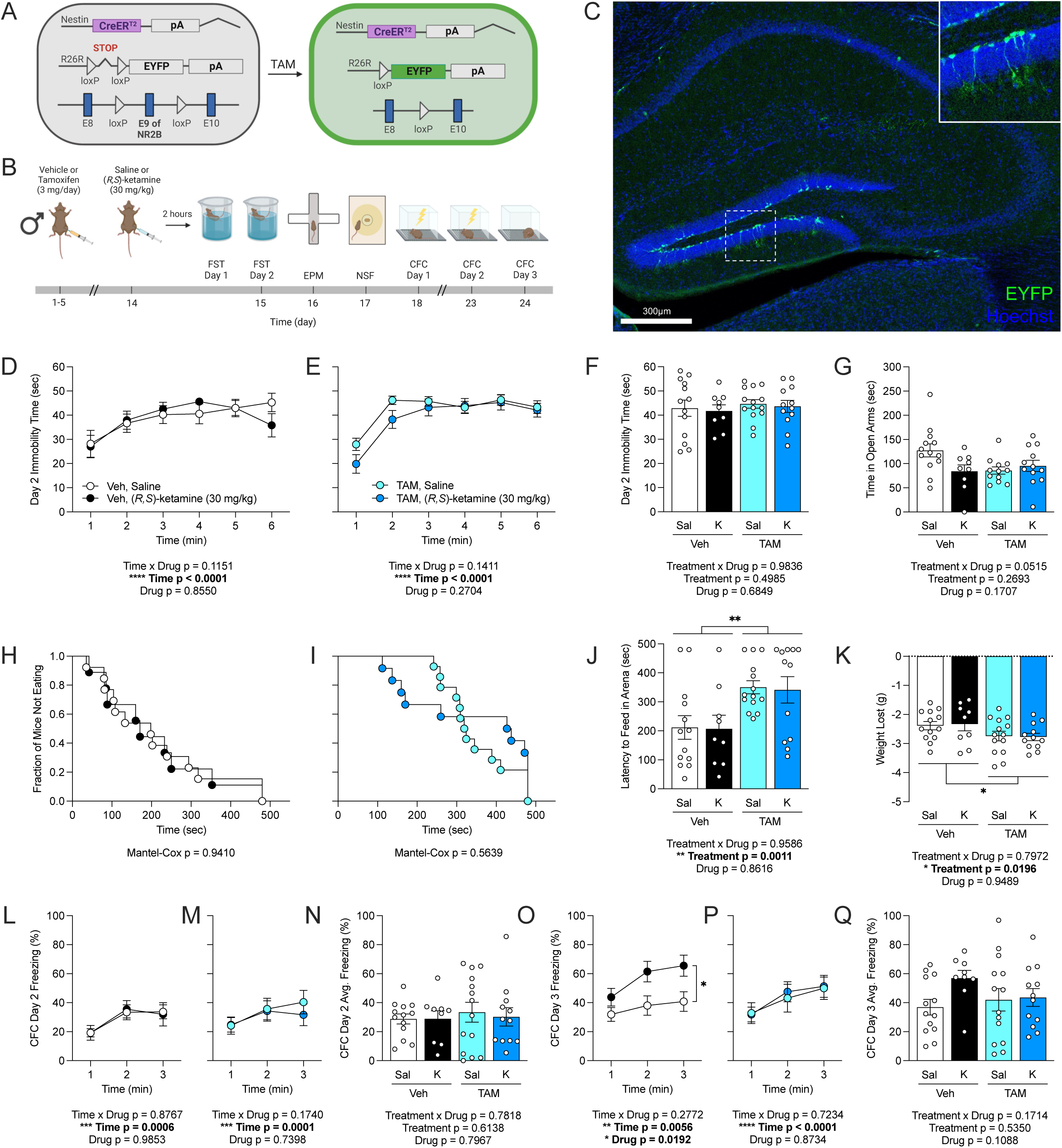
Deletion of the NR2B subunit from 2-week-old adult-born hippocampal neurons selectively influences (*R,S*)-ketamine’s fear effects in male mice. (**A**) Genetic schematic. (**B**) Experimental timeline. (**C**) Representative section showing EYFP (green) and Hoechst (blue) labeling in the HPC of NestinCreER^T2^ x NR2B x EYFP mice. (**D-F**) Immobility time on day 2 of the FST was comparable in all groups. (**G**) Time spent in the open arms of the EPM did not differ across groups. (**H-J**) In the NSF, there was an effect of Treatment on the latency to feed. TAM mice exhibited an increased latency when compared with Veh mice. (**K**) There was an effect of Treatment on weight loss in the NSF. (**L-N**) On day 2 of CFC, all groups exhibited comparable freezing. (**O**) On day 3 of CFC, K Veh mice froze significantly more than Sal Veh mice. (**P**) Both groups of TAM mice froze comparably. (**Q**) On day 3 of CFC, there was no impact of Treatment or Drug on average freezing behavior. (n = 8-14 male mice / group). Error bars represent ± SEM. * p < 0.05; ** p < 0.01; *** p < 0.001; **** p < 0.0001. HPC, hippocampus; EYFP, enhanced yellow fluorescent protein; Veh, vehicle; TAM, tamoxifen; FST, forced swim test; EPM, elevated plus maze; NSF, novelty suppressed feeding; CFC, contextual fear conditioning; sec, seconds; min, minutes; h, hours; mg, milligrams; kg, kilograms; g, grams; Sal, saline; K, (*R,S*)-ketamine (30 mg/kg).

On day 1 (**Fig. S3a-3c**) and day 2 of the FST (**Fig. 3d-3f**), and in the open (**Fig. 3g**) and closed arms (**Fig. S3d**) of the EPM, there was no significant effect of Treatment or Drug. In the NSF, there was an effect of Treatment on the latency to feed (**Fig. 3h-3j**) and on weight loss (**Fig. 3k**), but no post-hoc tests reached significance. On day 1 (**Fig. S3e-3g**) and day 2 (**Fig. 3l-3n**) of CFC, there was no effect of Drug or Treatment on freezing behavior. However, on day 3 of CFC, (*R*,*S*)-ketamine-injected Veh-treated mice had increased fear expression when compared with saline-injected Veh-treated mice (**Fig. 3o**). Notably, there was no effect of Drug in TAM-treated mice (**Fig. 3p-3q**). Overall, these data are in contrast with our 6-week-old experiment and suggest that TAM may be interfering with (*R,S*)-ketamine efficacy in control (Veh-injected) mice.

### Ablation of adult hippocampal neurogenesis does not influence (*R,S*)-ketamine efficacy

We next sought to extend our findings to a complete ablation of adult neurogenesis. Here, we utilized glial fibrillary acidic protein-thymidine kinase (GFAP-TK) mice, in which adult neurogenesis can be suppressed with ganciclovir (GCV) administration (**Fig. 4a**). Wild-type (WT) and GFAP-TK mice were implanted with mini-osmotic pumps filled with GCV for 4 weeks and 6 weeks later were administered a single injection of saline or (*R,S*)-ketamine (30 mg/kg) (**Fig. 4b**). GCV resulted in an ablation of neurogenesis in the GFAP-TK mice as indicated by the lack of doublecortin^+^ (DCX^+^) cells (**Fig. 4c**). Notably, these mice were maintained on a C57BL/6J background, in contrast to the Nestin line maintained on a 129S6/SvEv background.

**Fig. 4.**
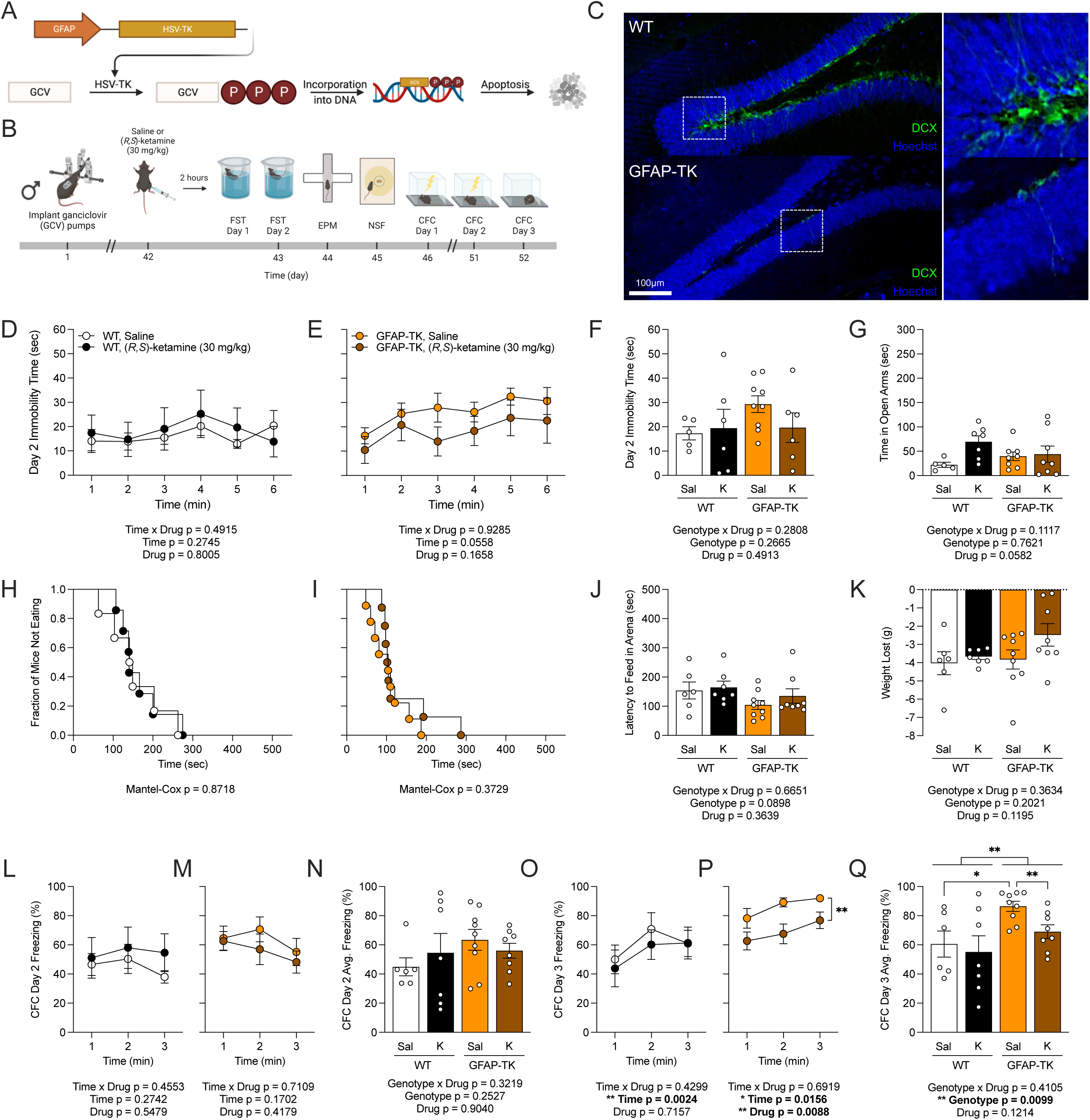
Ablation of adult hippocampal neurogenesis does not influence (*R,S*)-ketamine efficacy. (**A**) Genetic schematic. (**B**) Experimental timeline. (**C**) Representative section showing DCX (green) and Hoechst (blue) labeling in the DG of WT and GFAP-TK mice, indicating effective ablation of neurogenesis in GFAP-TK mice. (**D-F**) On day 2 of the FST, there was no effect of Genotype or Drug on immobility time. (**G**) Time in the open arms of the EPM did not differ across groups. (**H-J**) In the NSF, all groups had a comparable latency to feed. (**K**) Weight loss in the NSF did not differ across groups. (**L-N**) On day 2 of CFC, the percentage of freezing did not differ between the groups. (**O**) On day 3 of CFC, the percentage of freezing did not differ between WT Sal or WT K mice. (**P**) However, GFAP-TK K mice froze less than GFAP-TK Sal mice. (**Q**) There was an effect of Genotype on average freezing. (n = 6-9 male mice / group). Error bars represent ± SEM. *p < 0.05; ** p < 0.01. DCX, doublecortin; DG, dentate gyrus; WT, wild-type; GFAP-TK, glial fibrillary acidic protein-thymidine kinase; GCV, ganciclovir; FST, forced swim test; EPM, elevated plus maze; NSF, novelty suppressed feeding; CFC, contextual fear conditioning; sec, seconds; min, minutes; mg, milligrams; kg, kilograms; g, grams; Sal; saline; K, (*R,S*)-ketamine (30 mg/kg).

Neither Genotype nor Drug influenced behavior in the FST (**Fig. 4d-4f, Fig. S4a-4c**), the EPM (**Fig. 4g, Fig. S4d**), or the NSF (**Fig. 4h-4k**). However, there was an effect of Genotype on fear expression during day 1 of CFC (**Fig. S4e-4g**). While there was no effect of Genotype or Drug on fear expression during day 2 of CFC (**Fig. 4l-4n**), there was an effect of Genotype during day 3 of CFC. GFAP-TK mice displayed increased freezing as compared to wild-type mice (**Fig. 4o-4q**); while (*R,S*)-ketamine did not impact freezing in wild-type mice (**Fig. 4o**), (*R,S*)-ketamine decreased freezing in GFAP-TK mice (**Fig. 4p-4q**). However, (*R,S*)-ketamine’s effects may be significantly influenced by the background strain, and, thus, further studies of neurogenesis are needed in lines similar to the NestinCreER^T2^ line. These data suggest that in C57BL/6J mice, neurogenesis buffers against heightened fear expression in a manner that mimics (*R,S*)-ketamine and is in line with a previous study, which found that adult neurogenesis supports associative fear learning and protects against fear-induced anxiety [40].

### Ablation of NR2B from inhibitory interneurons does not influence (*R,S*)-ketamine efficacy in male mice

Next, to determine whether (*R,S*)-ketamine’s antidepressant effects require the NR2B subunit on inhibitory interneurons, which has previously been suggested [41, 42], we generated a triple transgenic mouse line that allowed for deletion of NR2B from parvalbumin^+^ (PV^+^) interneurons on a 129S6/SvEv background (**Fig. 5a**). In PV-Cre(-) and PV-Cre(+) mice, saline or (*R,S*)-ketamine (30 mg/kg) was administered at 14 weeks of age to match when Nestin mice were administered drug (**Fig. 5b**). Deletion of NR2B in PV^+^ cells was indicated by sparse EYFP labeling in the HPC (**Fig. 5c**).

**Fig. 5.**
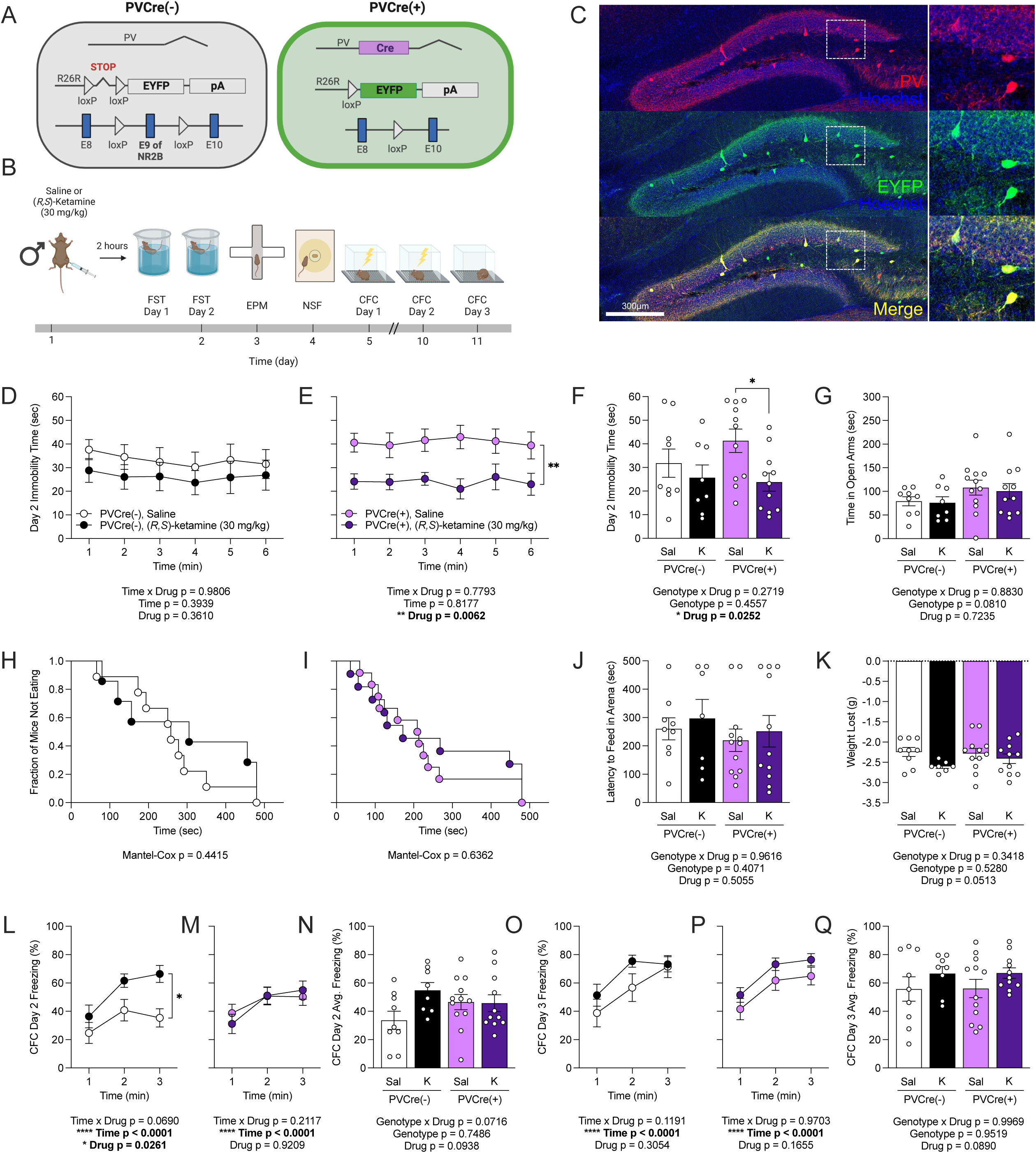
Ablation of NR2B from inhibitory interneurons does not influence (*R,S*)-ketamine efficacy in male mice. (**A**) Genetic schematic. (**B**) Experimental timeline. (**C**) Representative section showing PV (red), EYFP (green), and Hoechst (blue) labeling in the HPC of PV-Cre x NR2B x EYFP male mice, indicating PV-expressing cells with the NR2B subunit deleted. (**D**) In PV-Cre(-) mice, there was no effect of K on immobility time during the FST. (**E**) However, in PV-Cre(+) mice, K-treated mice had decreased immobility time in the FST as compared to Sal-treated mice. (**F**) K-administered PV-Cre(+) mice had decreased average immobility time in the FST than Sal-administered PV-Cre(+) mice. (**G**) Time in the open arms of the EPM did not differ across groups. (**H**) In the NSF, neither PV-Cre(-) nor (**I**) PV-Cre(+) mice differed in fraction of mice not eating over time in the NSF. (**J**) There was no difference in latency to feed or in (**K**) weight loss in the NSF across groups. (**L**) The percentage of freezing on day 2 of the CFC was increased in K-administered PV-Cre(-) mice as compared to Sal-administered PV-Cre(-) mice. (**M**) PV-Cre(+) Sal- or K-treated mice had comparable levels of freezing on day 2 of the CFC. (**N**) Average freezing on day 2 of the CFC did not differ across groups. (**O**) The percentage of freezing did not differ among PV-Cre(-) or (**P**) PV-Cre(+) Sal- or K-treated mice on CFC Day 3, a trend which was reflected in (**Q**) average freezing across groups. (n = 8-12 male mice / group). Error bars represent ± SEM. * p < 0.05; ** p < 0.01; **** p < 0.0001. PV, parvalbumin; EYFP, enhanced yellow fluorescent protein; HPC, hippocampus; FST, forced swim test; EPM, elevated plus maze; NSF, novelty suppressed feeding; CFC, contextual fear conditioning; sec, seconds; min, minutes; g, grams; K, (*R,S*)-ketamine (30 mg/kg).

There was an effect of Drug on day 1 (**Fig. S5a-5c**) and day 2 of the FST (**Fig. 5d-5f**). (*R*,*S*)-ketamine-injected PV-Cre(+) mice had significantly decreased immobility time as compared to saline-injected PV-Cre(+) mice (**Fig. 5e-5f**). There was no effect of Genotype or Drug on time in the open arms of the EPM (**Fig. 5g**), but there was an effect of Genotype on time in the closed arms (**Fig. S5d**). There was no effect of Genotype or Drug in the NSF (**Fig. 5h-5k**). Finally, in CFC, there was an effect of Drug on day 2 (**Fig. 5l-5n**), but no effect of Drug or Genotype on days 1 (**Fig. S5e-5g**) and 3 (**Fig. 5o-5q**). Intriguingly, these data indicate (*R*,*S*)-ketamine is effective against behavioral despair in experimental (i.e., PV-Cre(+)), but not in control mice (i.e., PV-Cre(-)).

### Ablation of NR2B from inhibitory interneurons influences (*R,S*)-ketamine’s effects on hyponeophagia in female mice

Finally, to validate our findings in male PV-Cre mice, saline or (*R,S*)-ketamine (10 mg/kg) was administered at 14 weeks of age to female PV-Cre(-) and PV-Cre(+) mice (**Fig. S6a-6b**). Deletion of NR2B in PV^+^ cells was indicated by sparse EYFP labeling in the HPC (**Fig. S6c**).

There was an effect of Drug and Genotype on day 1 of the FST (**Fig. S7a-7c**), with (*R*,*S*)-ketamine-injected PV-Cre(+) mice showing lower immobility time than saline-injected PV-Cre(+) mice (**Fig. S7b**). On FST day 2 (**Fig. S6d-6f**) and in the open (**Fig. S6g**) and closed (**Fig. S7d**) arms of the EPM, there was no effect of Drug or Genotype on respective behaviors. In the NSF, there was an effect of Drug (**Fig. S6h-6j**), where (*R,S*)-ketamine reduced feeding latency in PV-Cre(-) (**Fig. S6h**) but not PV-Cre(+) mice (**Fig. S6i**). There was no effect of Drug or Genotype on weight loss (**Fig. S6k**). Lastly, in CFC, (*R,S*)-ketamine significantly increased freezing in PV-Cre(-) (**Fig. S6l**) but not in PV-Cre(+) mice (**Fig. S6m-6n**) on day 2, though neither Genotype nor Drug affected behavior on day 1 (**Fig. S7e-7g**) and 3 (**Fig. S6o-6q**). Overall, these data indicate that NR2B deletion from PV^+^ cells in female mice mimics the effects of (*R*,*S*)-ketamine on hyponeophagia.

## DISCUSSION

Here, we have shown that NR2B expression on 6-week-old abGCs is necessary for (*R,S*)-ketamine’s effects on behavioral despair, hyponeophagia, and fear behavior in male mice, and for effects on hyponeophagia in female mice. To validate our findings in abGCs, we sought to determine the impact of NR2B expression on an interneuron PV^+^ population. Surprisingly, deletion of NR2B expression on PV^+^ interneurons produced divergent effects in males and females when (*R,S*)-ketamine was administered. In male mice, (*R*,*S*)-ketamine was effective at decreasing behavioral despair in PV-Cre(+) but not PV-Cre(-) mice. In female mice, NR2B deletion from PV^+^ cells mimics the effects of (*R*,*S*)-ketamine on hyponeophagia. Overall, these data highlight the importance of background strain, dosage, and sex for (*R,S*)-ketamine efficacy, and they suggest that further studies are warranted to reveal the precise mechanism by which the NR2B subunit in 6-week-old abGCs mediates (*R,S*)-ketamine’s effects.

In Veh-injected NestinCreER^T2^(+) x NR2B^f/f^ male mice, (*R,S*)-ketamine decreased behavioral despair, hyponeophagia, and fear expression, a finding that has been previously shown in 129S6/SvEv wild-type mice [7–9, 43]. These effects were occluded in TAM-injected NestinCreER^T2^(+) x NR2B^f/f^ mice lacking NR2B on 6-week-old abGCs. While the Nestin promoter is generally considered to label undifferentiated mitotic progenitor cells in the subventricular zone (SVZ) of the DG [44], it is important to note that the NestinCreER^T2^ line used in this study is one of several lines where leakiness is known to occur [45]. Indeed, EYFP expression in CA2 of the HPC was evident in TAM-treated mice. Nevertheless, this NestinCreER^T2^ line has limited ectopic expression overall [45], and these data are in accord with studies by Duman and colleagues showing NR2B is a key mediator in the antidepressant effects of (*R,S*)-ketamine [46, 47].

In female NestinCreER^T2^ mice, (*R,S*)-ketamine (10 mg/kg) was not as effective as an antidepressant as it was in male mice. However, (*R,S*)-ketamine alleviated hyponeophagia, which was occluded in TAM-injected NestinCreER^T2^(+) x NR2B^f/f^ mice. Our data align with a previous study using a viral strategy to ablate NR2B from somatostatin (SST)-expressing cortical neurons, which showed that in both male and female mice, (*R,S*)-ketamine (10 mg/kg) reduced hyponeophagia, and that knockdown of NR2B occluded this effect [41]. It is also possible that (*R,S*)-ketamine at this dose (10 mg/kg) alters the metabolism of female mice, as we also found that (*R,S*)-ketamine reduced weight in both Veh- and TAM-treated female but not male mice. We did observe scattered EYFP expression in the hypothalamus, which is involved in mediating metabolism, though this labeling was equally present in males and females (data not shown). However, we have previously shown that (*R,S*)-ketamine prevents stress-induced weight loss in males [6], revealing a novel sex-specific role of (*R,S*)-ketamine on body weight. Overall, these data highlight the need for further studies on sex differences in the behavioral and genetic phenotypes of MDD and whether different dose concentrations of (*R,S*)-ketamine target similar neural and molecular pathways across both sexes.

Deletion of NR2B from up to 2-week-old Nestin^+^ abGCs unexpectedly led to an increase in the latency to feed in the NSF regardless of (*R,S*)-ketamine treatment. Surprisingly, at this 2-week timepoint, (*R,S*)-ketamine was not effective at decreasing behavioral despair, hyponeophagia, or fear behavior in Veh-treated NestinCreER^T2^(+) x NR2B^f/f^ mice, but TAM-treated mice exhibited hyponeophagia and lost more weight as compared to Veh-treated mice. It has previously been shown that TAM injection can induce long-term toxicity in mice; increased Cre induction has been observed up to 4 weeks following the start of TAM dosing, which can be due to gradual release of TAM from the vehicle and the stability of TAM *in vivo* [48]. Here, mice were administered 5 doses of Veh or TAM and tested only 2 weeks later, which may have altered baseline behavior and occluded (*R,S*)-ketamine’s antidepressant effects. Although TAM was chosen for this study, it is possible that utilization of its metabolite 4-hydroxytamoxifen (4-OHT), which has a shorter half-life [49], may be more advantageous for determining the age of cells that may be mediating (*R,S*)-ketamine’s antidepressant effects. Thus, future studies should consider the use of 4-OHT for Cre inducible lines to assess behavior more proximal to the drug injections.

To determine whether the antidepressant effects of (*R,S*)-ketamine are specific to NR2B-containing 6-week-old abGCs, we assessed whether all abGCs are necessary by fully ablating neurogenesis. While prior studies have shown that neurogenesis supports contextual fear memory [35, 50], we found in current experiments that GFAP-TK-mediated ablation of neurogenesis did not impair CFC performance. However, the mice in this study underwent several stressful behavioral paradigms (FST, EPM, NSF) prior to CFC, whereas previous experiments isolated the effects of neurogenesis ablation by minimizing aversive or stressful stimuli [50]. Indeed, our current results are more consistent with evidence that neurogenesis buffers against the effects of stress [40, 51], including heightened freezing behavior, as seen with ablation of neurogenesis in the present study. However, treatment with (*R,S*)-ketamine rescued this heightened fear, suggesting that it retains the ability to buffer against stress-enhanced fear even in the absence of adult neurogenesis. Prior data also suggest that (*R,S*)-ketamine activates abGCs [32] and causes them to rapidly mature [52], so this effect may have been mediated through a small population of recently-born abGCs that escaped GFAP-TK ablation. These data complement previous data from our lab showing ventral CA3 (vCA3) is necessary (*R,S*)-ketamine efficacy [43], suggesting that (*R,S*)-ketamine may also directly activate vCA3 targets to buffer against stress-enhanced fear even when adult neurogenesis is suppressed. Nonetheless, behavioral effects of treatment were absent in more traditional assays of anxiety- and depression-like behavior, consistent with the idea that (*R,S*)-ketamine targets abGCs for most antidepressant effects. Finally, these mice were also on a C57BL/6J background, which we have shown to be insensitive to antidepressant (*R,S*)-ketamine unless administered after chronic corticosterone (CORT) treatment [6]. Overall, these data indicate that neurogenesis ablation increases stress-enhanced fear expression, and that (*R,S*)-ketamine can effectively buffer this fear, possibly by inducing the rapid maturation and activation of abGCs.

Finally, we tested if NR2B expression on PV interneurons was necessary for (*R,S*)-ketamine efficacy. In male mice, (*R*,*S*)-ketamine was surprisingly effective at decreasing behavioral despair in the experimental PV-Cre(+) but not control PV-Cre(-) x NR2B^f/f^ mice. In female mice, (*R*,*S*)-ketamine was effective at decreasing behavioral despair on day 1 but not day 2 of the FST in PV-Cre(+) but not PV-Cre(-) x NR2B^f/f^ mice. There are several reasons that may explain the discrepancy between this current study and prior studies, as well as with other lines included in this study. Firstly, PVCre mice are on a full 129S6/SvEv background, whereas NestinCreER^T2^ mice are mixed 129S6/SvEv and C57BL/6J. We have previously shown that (*R*,*S*)-ketamine (30 mg/kg) is ineffective as an antidepressant in non-stressed 129S6/SvEv mice [6]. Secondly, NestinCreER^T2^ mice underwent the stress of multiple daily injections 6 weeks prior to testing, whereas the PVCre mice are a constitutive deletion line that does not require TAM injections. This genetic strategy may have influenced responsiveness to (*R,S*)-ketamine. Thirdly, we did not use a viral/genetic strategy in this study as the Duman group used, which limited NR2B deletion to just the PFC [41] as opposed to all PV interneurons. Future studies selectively deleting NR2B from hippocampal interneurons using viral strategies will be necessary. Finally, Duman and colleagues report that (*R*,*S*)-ketamine’s effects are blocked in PVCre/adeno-associated virus (AAV) mice, but they also show that genetic deletion alters baseline behavior, as these mice exhibit significantly decreased immobility compared to WT/AAV mice [41]. Our data are in accord with this finding in that the genetically deleted line shows significantly altered baseline behavior.

In summary, these experiments demonstrate a putative target for (*R,S*)-ketamine’s rapid-acting antidepressant effects. Because we have also previously found a protective effect of (*R,S*)-ketamine against stress-induced pathophysiology [6–12], it will be critical to determine whether the mechanisms of (*R,S*)-ketamine’s antidepressant and prophylactic effects overlap or diverge. These data will inform future studies to identify specific populations of cells that mediate antidepressant effects and that may be targeted via more rapid-acting, efficacious therapeutics.

## ACKNOWLEDGMENTS

We thank members of the Denny Laboratory for their insightful comments on this project and manuscript. We also thank Dr. René Hen for valuable input on this project and for the use of transgenic mice.

## AUTHOR CONTRIBUTIONS

NEB, JCM, SCL, MRD, and CAD contributed to the conception and design of the work, the analysis and interpretation of the data for the work, drafting the work, and revising it critically for important intellectual content. NEB, JCM, LRL, CTL, SCL, CXS, RAB, ES, IP, and DS contributed to the acquisition and presentation of the data. NEB, JCM, and CAD approved the version of the manuscript to be published and agreed to be accountable for all aspects of the work in ensuring that questions related to the accuracy and integrity of any part of the work are appropriately investigated and resolved. All authors contributed to the article and approved the submitted version.

## ADDITIONAL INFORMATION

**Supplementary information:** The online version contains supplementary material available at [TBD].

**Correspondence** and requests for materials should be addressed to Christine Ann Denny.

## DATA AVAILABILITY STATEMENT

The raw data supporting the conclusions of this article will be made available by the authors, without undue reservation.

## ETHICS STATEMENT

All experiments were approved by the Institutional Animal Care and Use Committees (IACUCs) at Columbia University Irving Medical Center (CUIMC), the New York State Psychiatric Institute (NYSPI), and the University of Texas at Austin.

## REFERENCES

1. Kessler RC, Chiu WT, Demler O, Merikangas KR, Walters EE. Prevalence, severity, and comorbidity of 12-month DSM-IV disorders in the National Comorbidity Survey Replication. Arch Gen Psychiatry. 2005;62:617–27.

2. World Health Organization. Depression. 2021. https://www.who.int/news-room/fact-sheets/detail/depression. Accessed 2 February 2023.

3. Mathers CD, Loncar D. Projections of global mortality and burden of disease from 2002 to 2030. PLoS Med. 2006;3:e442.

4. Sackeim HA. The definition and meaning of treatment-resistant depression. J Clin Psychiatry. 2001;62:10–7.

5. Autry, AE, Adachi M, Nosyreva E, Na ES, Los MF, Cheng P, et al. NMDA receptor blockade at rest triggers rapid behavioural antidepressant responses. Nature. 2011;475:91–5.

6. Brachman, RA, McGowan JC, Perusini JN, Lim SC, Plam TH, Faye C, et al. Ketamine as a prophylactic against stress-induced depressive-like behavior. Biol Psychiatry. 2016;79:776– 86.

7. McGowan JC, LaGamma CT, Lim SC, Tsitsiklis M, Neria Y, Brachman RA, et al. Prophylactic ketamine attenuates learned fear. Neuropsychopharmacology. 2017;42:1577– 89.

8. McGowan JC, Hill C, Mastrodonato A, LaGamma CT, Kitayev A, Brachman RA, et al. Prophylactic ketamine alters nucleotide and neurotransmitter metabolism in brain and plasma following stress. Neuropsychopharmacology. 2018;43:1813–21.

9. Mastrodonato, A, Cohensedgh, O, LaGamma, CT, McGowan, JC, Hunsberger, HC, Denny, CA. Prophylactic (R,S)-ketamine selectively protects against inflammatory stressors. Behav Brain Res. 2020;378:112238.

10. Chen BK, Mendez-David I, Luna VM, Faye C, Gardier AM, David DJ, et al. Prophylactic efficacy of 5-HT4R agonists against stress. Neuropsychopharmacology. 2020;45:542–55.

11. Chen BK, Luna VM, LaGamma CT, Xu X, Deng SX, Suckow RF, et al. Sex-specific neurobiological actions of prophylactic (R,S)-ketamine, (2R,6R)-hydroxynorketamine, and (2S,6S)-hydroxynorketamine. Neuropsychopharmacology. 2020;45:1545–56.

12. Chen BK, Luna VM, Shannon ME, Hunsberger HC, Mastrodonato AM, Stackmann M, et al. Fluoroethylnormemantine, a novel NMDA receptor antagonist, for the prevention and treatment of stress-induced maladaptive behavior. Biol Psychiatry. 2021;90:458–72.

13. Krystal JH, Karper LP, Seibyl JP, Freeman GK, Delaney R, Bremner JD, et al. Subanesthetic effects of the noncompetitive NMDA antagonist, ketamine, in humans. Psychotomimetic, perceptual, cognitive, and neuroendocrine responses. Arch Gen Psychiatry. 1994;51:199–214.

14. Mathews DC, Henter ID, Zarate CA. Targeting the glutamatergic system to treat major depressive disorder: rationale and progress to date. Drugs. 2012;72:1313–33.

15. Skolnick P, Layer RT, Popik P, Nowak G, Paul IA, Trullas R. Adaptation of N-methyl-D-aspartate (NMDA) receptors following antidepressant treatment: implications for the pharmacotherapy of depression. Pharmacopsychiatry. 1996;29:23–6.

16. Monyer H, Sprengel R, Schoepfer R, Herb A, Higuchi M, Lomeli H, et al. Heteromeric NMDA receptors: molecular and functional distinction of subtypes. Science. 1992;256:1217–21.

17. Laube B, Hirai H, Sturgess M, Betz H, Kuhse J. Molecular determinants of agonist discrimination by NMDA receptor subunits: analysis of the glutamate binding site on the NR2B subunit. Neuron. 1997;18:493–503.

18. Feyissa AM, Chandran A, Stockmeier CA, Karolewicz B. Reduced levels of NR2A and NR2B subunits of NMDA receptor and PSD-95 in the prefrontal cortex in major depression. Prog Neuropsychopharmacol Biol Psychiatry. 2009;33:70–5.

19. Saez E, Erkoreka L, Moreno-Calle T, Berjano B, Gonzalez-Pinto A, Basterreche N, et al. Genetic variables of the glutamatergic system associated with treatment-resistant depression: A review of the literature. World J Psychiatry. 2022;12:884–96.

20. Miller OH, Yang L, Wang CC, Hargroder EA, Zhang Y, Delpire E, et al. GluN2B-containing NMDA receptors regulate depression-like behavior and are critical for the rapid antidepressant actions of ketamine. Elife. 2014;3:e03581.

21. Preskorn SH, Baker B, Kolluri S, Menniti FS, Krams M, Landen JW. An innovative design to establish proof of concept of the antidepressant effects of the NR2B subunit selective N-methyl-D-aspartate antagonist, CP-101,606, in patients with treatment-refractory major depressive disorder. J Clin Psychopharmacol. 2008;28:631–7.

22. Lima-Ojeda JM, Vogt MA, Pfeiffer N, Dormann C, Köhr G, Sprengel R, et al. Pharmacological blockade of GluN2B-containing NMDA receptors induces antidepressant-like effects lacking psychotomimetic action and neurotoxicity in the perinatal and adult rodent brain. Prog Neuropsychopharmacol Biol Psychiatry. 2013;45:28–33.

23. Stasiuk W, Szopa A, Serefko A, Wyska E, Świąder K, Dudka J, et al. Influence of the selective antagonist of the NR2B subunit of the NMDA receptor, traxoprodil, on the antidepressant-like activity of desipramine, paroxetine, milnacipran, and bupropion in mice. J Neural Transm. 2017;124:387–96.

24. Ibrahim L, Diaz Granados N, Jolkovsky L, Brutsche N, Luckenbaugh DA, Herring WJ, et al. A Randomized, placebo-controlled, crossover pilot trial of the oral selective NR2B antagonist MK-0657 in patients with treatment-resistant major depressive disorder. J Clin Psychopharmacol. 2012;32:551–7.

25. Duman RS. Ketamine and rapid-acting antidepressants: a new era in the battle against depression and suicide [version 1; peer review: 3 approved]. F1000 Fac Rev. 2018;7:659.

26. Boldrini M, Underwood MD, Hen R, Rosoklija GB, Dwork AJ, Mann JJ, et al. Antidepressants increase progenitor cells in the human hippocampus. Neuropsychopharmacology. 2009;34:2376–89.

27. Santarelli L, Saxe M, Gross C, Surget A, Battaglia F, Dulawa S, et al. Requirement of hippocampal neurogenesis for the behavioral effects of antidepressants. Science. 2003;301:805–9.

28. Perera TD, Dwork AJ, Keegan KA, Thirumangalakudi L, Lipira CM, Joyce N, et al. Necessity of hippocampal neurogenesis for the therapeutic action of antidepressants in adult nonhuman primates. PLoS One. 2011;6:e17600.

29. Yamada J, Jinno S. Potential link between antidepressant-like effects of ketamine and promotion of adult neurogenesis in the ventral hippocampus of mice. Neuropharmacology. 2019;158:107710.

30. Planchez B, Lagunas N, Le Guisquet A-M, Legrand M, Surget A, Hen R, et al. Increasing adult hippocampal neurogenesis promotes resilience in a mouse model of depression. Cells. 2021;10:972.

31. Clarke M, Razmjou S, Prowse N, Dwyer Z, Litteljohn D, Pentz R, et al. Ketamine modulates hippocampal neurogenesis and pro-inflammatory cytokines but not stressor induced neurochemical changes. Neuropharmacology. 2017;112:210–20.

32. Rawat R, Tunc-Ozcan E, McGuire TL, Peng C-Y, Kessler JA. Ketamine activates adult-born immature granule neurons to rapidly alleviate depression-like behaviors in mice. Nat Commun. 2022;13:2650.

33. Schmidt-Hieber C, Jonas P, Bischofberger J. Enhanced synaptic plasticity in newly generated granule cells of the adult hippocampus. Nature. 2004;429:184–7.

34. Sahay A, Scobie KN, Hill AS, O’Carroll CM, Kheirbek MA, Burghardt NS, et al. Increasing adult hippocampal neurogenesis is sufficient to improve pattern separation. Nature. 2011;472:466–70.

35. Denny CA, Burghardt NS, Schachter DM, Hen R, Drew MR. 4- to 6-week-old adult-born hippocampal neurons influence novelty-evoked exploration and contextual fear conditioning. Hippocampus. 2012;22:1188–201.

36. Burghardt NS, Park EH, Hen R, Fenton AA. Adult-born hippocampal neurons promote cognitive flexibility in mice. Hippocampus. 2012;22:1795–1808.

37. Hill AS, Sahay A, Hen R. Increasing adult hippocampal neurogenesis is sufficient to reduce anxiety and depression-like behaviors. Neuropsychopharmacology. 2015;40:2368–78.

38. Huckleberry KA, Shue F, Copeland T, Chitwood RA, Yin W, Drew MR. Dorsal and ventral hippocampal adult-born neurons contribute to context fear memory. Neuropsychopharmacology. 2018;43:2487–96.

39. Chen BK, Denny CA. Weapons of stress reduction: (*R*,S)-ketamine and its metabolites as prophylactics for the prevention of stress-induced psychiatric disorders. Neuropharmacology. 2023;224:10934.

40. Seo D-O, Carillo MA, Lim SC-H, Tanaka KF, Drew MR. Adult hippocampal neurogenesis modulates fear learning through associative and nonassociative mechanisms. J Neurosci. 2015;35:11330–45.

41. Gerhard DM, Pothula S, Liu RJ, Wu M, Li X-Y, Girgenti MJ, et al. GABA interneurons are the cellular trigger for ketamine’s rapid antidepressant actions. J Clin Invest. 2020;130:1336–49.

42. Ng LHL, Huang Y, Han L, Chang RC-C, Chan YS, Lai CSW. Ketamine and selective activation of parvalbumin interneurons inhibit stress-induced dendritic spine elimination. Transl Psychiatry. 2018;8:272.

43. Mastrodonato A, Martinez R, Pavlova IP, LaGamma CT, Brachman RA, Robison AJ, et al. Ventral CA3 activation mediates prophylactic ketamine efficacy against stress-induced depressive-like behavior. Biol Psychiatry. 2018;84:846–56.

44. Fukuda S, Kato F, Tozuka Y, Yamaguchi M, Miyamoto Y, Hisatsune T. Two distinct subpopulations of nestin-positive cells in adult mouse dentate gyrus. J Neurosci. 2003;23:9357–66.

45. Sun MY, Yetman MJ, Lee TC, Chen Y, Jankowsky JL. Specificity and efficiency of reporter expression in adult neural progenitors vary substantially among nestin-CreER(T2) lines. J Comp Neurol. 2014;522:1191–208.

46. Li N, Lee B, Liu R-J, Banasr M, Dwyer JM, Iwata M, et al. mTOR-dependent synapse formation underlies the rapid antidepressant effects of NMDA antagonists. Science. 2010;329:959–64.

47. Ali F, Gerhard DM, Sweasy K, Pothula S, Pittenger C, Duman RS, et al. Ketamine disinhibits dendrites and enhances calcium signals in prefrontal dendritic spines. Nat Commun. 2020;11:72.

48. Donocoff RS, Teteloshvili N, Chung H, Shoulson R, Creusot RJ. Optimization of tamoxifen-induced Cre activity and its effect on immune cell populations. Sci Rep. 2020;10:1–12.

49. Cazzulino AS, Martinez R, Tomm NK, Denny CA. Improved specificity of hippocampal memory trace labeling. Hippocampus. 2016;26:752–62.

50. Drew MR, Denny CA, Hen R. Arrest of adult hippocampal neurogenesis in mice impairs single-but not multiple-trial contextual fear conditioning. Behav Neurosci. 2010;124:446–54.

51. Snyder JS, Soumier A, Brewer M, Pickel J, Cameron HA. Adult hippocampal neurogenesis buffers stress responses and depressive behaviour. Nature 2011;476:458–61.

52. Soumier A, Carter RM, Schoenfeld TJ, Cameron HA. New hippocampal neurons mature rapidly in response to ketamine but are not required for its acute antidepressant effects on neophagia in rats. eNeuro. 2016;3:ENEURO.0116-5.2016.

