## Supplemental Information for "(*R,S*)-ketamine’s rapid-acting antidepressant effects are modulated by NR2B-containing NMDA receptors on adult-born hippocampal neurons"

### SUPPLEMENTAL METHODS

#### Mice

All mice were generated and bred in-house with genotypes confirmed by published PCR protocols [1–3].

*NestinCreER$^{T2}$ x NR2B x enhanced yellow fluorescent protein (EYFP) mice*: NestinCreER$^{T2}$ [4], homozygous NR2B exon 9-floxed (f/f) [3], and homozygous R26R-STOP-floxed (f/f) EYFP transgenic mice [5] were generated as previously described [4, 6]. Experimental mice were generated by breeding NestinCreER$^{T2}$, NR2B$^{f/f}$, EYFP$^{f/f}$ with NR2B$^{f/f}$, EYFP$^{f/f}$ mice. Male and female mice backcrossed to a mostly 129S6/SvEv background were used for experiments as specified in the text.

*Parvalbumin (PV)-Cre x NR2B x EYFP mice:* PV-Cre mice were generated as previously described [7]. Experimental male and female mice were generated by breeding PV-Cre, NR2B$^{f/f}$, EYFP$^{f/f}$ [5] with NR2B$^{f/f}$, EYFP$^{f/f}$ mice and backcrossed to a 129S6/SvEv background for maintenance.

*Glial fibrillary acidic protein-thymidine kinase (GFAP-TK) mice:* Glial fibrillary acidic protein-thymidine kinase (GFAP-TK) mice were generated [8, 9] and backcrossed to a C57BL/6J background for maintenance.

#### Drugs

*Ganciclovir (GCV)*

GCV was administered to wild-type (WT) and GFAP-TK mice for 4 weeks via subcutaneous osmotic minipumps (Alzet Model 1004, 0.11μL/h). The timing of GCV administration with respect to behavioral testing is described in the main text. Briefly, pumps were first filled with

60mg/mL GCV in sterile 0.9% saline. Mice were anesthetized with isoflurane (2-4%). A small incision was made on the upper dorsum. A hemostat was inserted to open a subcutaneous pocket for the osmotic minipump, which was inserted parasagitally on the back. The incision was closed with sutures. To prevent necrosis at the drug infusion site, the pumps were gently rotated within the subcutaneous space every other day.

**Behavioral Assays**

*Forced Swim Test (FST)*

The FST is typically used in rodents to screen for antidepressant activity [10, 11]. In the FST, time spent immobile, as opposed to swimming, is used as a measure of behavioral despair. Mice were administered the FST as previously described [12]. Briefly, mice were placed into clear plastic buckets 20 cm in diameter and 23 cm deep filled 2/3 of the way with 22°C water. Mice were videotaped from the side for 6 min and were exposed to the swim test on 2 consecutive days. Scoring was performed using an automated Viewpoint Videotrack software package (http://www.viewpoint.fr/en/a/anxiety-and-depression).

*Elevated Plus Maze (EPM)*

Testing was performed as previously described [13]. Briefly, the maze is a plus-cross-shaped apparatus consisting of four arms, two brightly lit open (800-900 lux) and two enclosed by walls, linked by a central platform at the height of 50 cm from the floor. Mice were individually placed in the center of the maze facing an open arm and were allowed to explore the maze for 5 min. The time spent in the open arms and/or closed arms was quantified. Videos were scored using ANY-maze behavior tracking software (Stoelting, Wood Dale, IL).

*Novelty Suppressed Feeding (NSF) Paradigm*

The NSF paradigm was performed as previously described [14]. Briefly, the testing apparatus consisted of a plastic box (50 x 50 x 20 cm). The floor was covered with approximately 2 cm of wooden bedding and the arena was brightly lit (1100-1200 lux). Mice were food restricted for 18 h. All food was removed from the home cage. At the time of testing, a single pellet of food (regular chow) was placed on a white paper platform positioned in the center of the box. Each animal was placed in a corner of the box, and a stopwatch was immediately started. The latency of the mice to begin eating was recorded. Immediately after the latency was recorded, the food pellet was removed from the arena. The mice were then assessed for post-restriction weight. Male and female mice were not run at the same time to avoid potential confounds. A Kaplan-Meier survival analysis was used due to the lack of normal distribution of data. The Mantel-Cox log-rank test was used to evaluate differences between the experimental groups.

*Contextual Fear Conditioning (CFC)*

CFC took place in Coulbourn fear conditioning boxes that contained one clear Plexiglas wall, three aluminum walls, and a stainless-steel grid floor as previously described [2, 9, 15]. In the two-trial contextual fear conditioning, the training context included a house light and fan, and lemon scent was placed under the grid floor. After 180 s, mice received a single 2 s foot shock of 0.75 mA. Mice were taken out 15 s after termination of the foot shock and returned to their home cage. The box was cleaned with 70% EtOH between runs. Mice received a second shock in the training context 5 days following the training session and were tested for conditioned fear during this exposure and during a second exposure 24 h later. The testing procedure and context of these two exposures were identical to those used on the training day with the exception that on the second exposure, a shock was not given. Both sessions were scored for freezing. Digital video cameras recorded the session; FreezeFrame and FreezeView software v4 (Actimetrics, Wilmette, IL) were used for recording and analyzing freezing behavior.

**Brain Processing and Immunohistochemistry**

Immunohistochemistry was performed as previously described [4, 16, 17]. Mice were perfused and brains were placed in 4% paraformaldehyde (PFA) in 1X phosphate buffered saline (PBS) overnight then subsequently in 1X PBS. At least 3 days later, brains were cut at 50 μm thickness on a vibratome (Leica Biosystems, Wetzlar, Germany) and stored in 1X PBS with 0.1% sodium azide.

All sections were washed in 1X PBS in 3 increments of 10 min each and then were blocked in 10% normal donkey serum (NDS) in 1X PBS with 0.1% Triton X-100 (0.1% PBST) for 2 hours at room temperature (RT). Incubation with primary antibody was performed at 4ºC overnight. For NestinCreER$^{T2}$ mice, sections were stained with chicken anti-green fluorescent protein (GFP) (1:500, ab13970, Abcam, Cambridge, United Kingdom) [4]. For PV-Cre mice, sections were stained with chicken anti-GFP (1:500, ab13970, Abcam, Cambridge, United Kingdom) and rabbit anti-PV (1:3,000, PV 27, Swant, Burgdorf, Switzerland) [16]. For GFAP-TK mice, sections were washed in 1X PBS in 3 increments of 10 min each and then were blocked in 5% NDS in 1X PBS with 0.25% Triton X-100 (0.25% PBST) for 1 hour at RT. Incubation at primary antibody was performed at 4ºC overnight. Sections were stained with rabbit anti-DCX (1:4,000, ab18723, Abcam, Cambridge, United Kingdom) [17].

After primary incubation, sections were washed 3 times in 1X PBS and incubated in secondary antibody overnight at 4ºC. For NestinCreER$^{T2}$ mice, sections were stained with donkey anti-chicken Cy2 (1:500, 703-225-155, Jackson ImmunoResearch, West Grove, PA) and Hoechst 33342 (1:10,000, Thermo Fisher Scientific, Waltham, MA). For PV-Cre mice, sections were stained with donkey anti-chicken Cy2 (1:500, 703-225-155, Jackson ImmunoResearch, West Grove, PA), donkey anti-rabbit Cy3 (1:500, 711-165-152, Jackson ImmunoResearch, West Grove, PA), and Hoechst 33342 (1:10,000, Thermo Fisher Scientific, Waltham, MA). For GFAP-TK mice, sections were stained with donkey anti-rabbit Alexa Fluor 488 (1:250, 711-545-152, Jackson ImmunoResearch, West Grove, PA) and Hoechst 33342 (1:10,000, Thermo Fisher Scientific, Waltham, MA). Following secondary antibody incubation, sections were then washed again 3 times in 1X PBS. Sections were mounted on slides and allowed to dry for approximately 30 min before adding mounting medium Fluoromount G (Electron Microscopy Sciences, Hatfield, PA) and a coverslip.

**Confocal Microscopy**

All samples were imaged on a confocal scanning microscope (Leica TCS SP8, Leica Microsystems Inc., Wetzlar, Germany) with 2 simultaneous PMT detectors, as previously described [18]. Cy2 and Alexa Fluor 488 were excited at 488 nm and detected at 500-550 nm, Cy3 was excited at 555 nm and detected at 577-627 nm, and Hoechst 33342 was excited with ultraviolet light at 350 nm and detected at 440-520 nm. Sections were imaged with a dry Leica 20x objective (NA 0.70, working distance 0.5 mm), with a pixel size of 1.08 x 1.08 $\mu m^2$, a z step of 1.5 μm, and a z-stack of 11 μm. Fields of view were stitched together to form tiled images by using an automated stage and tiling function and algorithm of the LAS X software. Images are represented as maximum-intensity projection of the z-stacks.

### SUPPLEMENTARY REFERENCES


1. Feil R, Wagner J, Metzger D, Chambon P. Regulation of cre recombinase activity by mutated estrogen receptor ligand-binding domains. Biochem Biophys Res Commun. 1997;237:752–7.

2. Denny CA, Kheirbek MA, Alba EL, Tanaka KF, Brachman RA, Laughman KB, et al. Hippocampal memory traces are differentially modulated by experience, time, and adult neurogenesis. Neuron. 2014;83:189–201.

3. Von Engelhardt J, Doganci B, Jensen V, Hvalby O, Gongrich C, Taylor A, et al. Contribution of hippocampal and extra-hippocampal NR2B-containing NMDA receptors to performance on spatial learning tasks. Neuron. 2008;60:846–60.

4. Dranovsky A, Picchini AM, Moadel T, Sisti AC, Yamada A, Kimura S, et al. Experience dictates stem cell fate in the adult hippocampus. Neuron. 2011;70:908–23.

5. Srinivas S, Watanabe T, Lin CS, William CM, Tanabe Y, Jessell TM, et al. Cre reporter strains produced by targeted insertion of EYFP and ECFP into the ROSA26 locus. BMC Dev Biol. 2001;1:4.

6. Kheirbek MA, Tannenholz L, Hen R. NR2B-dependent plasticity of adult-born granule cells is necessary for context discrimination. J Neurosci. 2012;32:8696–702.

7. Hippenmeyer S, Vrieseling E, Sigrist M, Portmann T, Laengle C, Ladle DR, et al. A developmental switch in the response of DRG neurons to ETS transcription factor signaling. PLoS Biol. 2005;3:e159.

8. Snyder JS, Soumier A, Brewer M, Pickel J, Cameron HA. Adult hippocampal neurogenesis buffers stress responses and depressive behavior. Nature. 2011;476:458–61.

9. Denny CA, Burghardt NS, Schachter DM, Hen R, Drew MR. 4- to 6-week-old adult-born hippocampal neurons influence novelty-evoked exploration and contextual fear conditioning. Hippocampus. 2012;22:1188–201.

10. Holmes PV. Rodent models of depression: reexamining validity without anthropomorphic inference. Crit Rev Neurobiol. 2003;15:143–74.

11. Petit-Demouliere B, Chenu F, Bourin M. Forced swimming test in mice: A review of antidepressant activity. Psychopharmacology. 2005;177:245–55.

12. Richardson-Jones JW, Craige CP, Guiard BP, Stephen A, Metzger KL, Kung HF, et al. 5-HT1A autoreceptor levels determine vulnerability to stress and response to antidepressants. Neuron. 2010;65:40–52.

13. Saxe MD, Battaglia F, Wang J-W, Malleret G, David DJ, Monckton JE, et al. Ablation of Hippocampal Neurogenesis Impairs Contextual Fear Conditioning and Synaptic Plasticity in the Dentate Gyrus. Proc Natl Acad Sci U S A. 2006;103:17501–6.

14. David DJ, Samuels BA, Riner Q, Wang JW, Marstellar D, Menez I, et al. Neurogenesis-dependent and -independent effects of fluoxetine in an animal model of anxiety/depression. Neuron. 2009;62:479-93.

15. Drew MR, Denny CA, Hen R. Arrest of adult hippocampal neurogenesis in mice impairs single-but not multiple-trial contextual fear conditioning. Behav Neurosci. 2010;124:446–54.

16. Chen SX, Kim AN, Peters AJ, Komiyama T. Subtype-specific plasticity of inhibitory circuits in motor cortex during motor learning. Nat Neurosci. 2015;18:1109–15.

17. Seo D-O, Carillo MA, Lim SC-H, Tanaka KF, Drew MR. Adult hippocampal neurogenesis modulates fear learning through associative and nonassociative mechanisms. J Neurosci. 2015;35:11330–45.

18. Pavlova IP, Shipley SC, Lanio M, Hen R, Denny CA. Comparison and optimization of whole-brain immunolabeling techniques for indelibly-labeled memory traces. Hippocampus. 2018;28:523–35.

### SUPPLEMENTARY FIGURES AND FIGURE LEGENDS

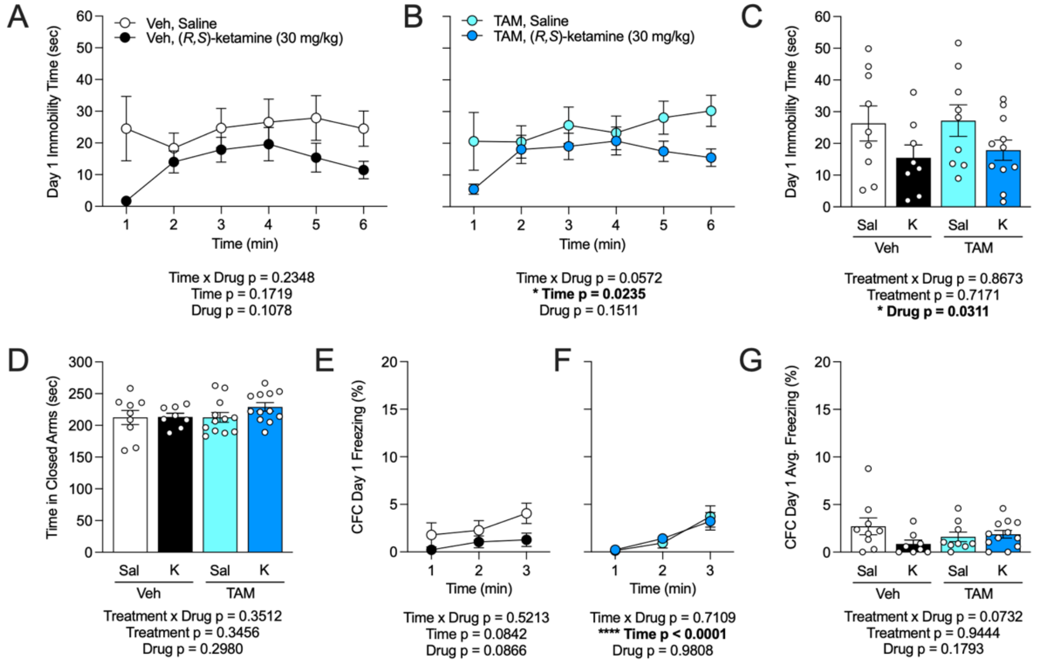


**Supplementary Fig. 1. Ablation of NR2B from 6-week-old adult-born neurons does not alter (*R,S*)-ketamine efficacy in male mice**. (**A**) On day 1 of the FST, there was no difference in immobility time between Veh-Sal and Veh-K mice or (**B**) between TAM-Sal and TAM-K mice. (**C**) While there was a main effect of Drug on average immobility time on day 1 of the FST, immobility did not differ between Veh- and TAM-treated groups. (**D**) Time spent in the closed arms of the EPM did not differ across groups. (**E**) Freezing on day 1 of the CFC did not differ between Veh-Sal and Veh-K or (**F**) between TAM-Sal and TAM-K mice. (**G**) Average freezing during CFC Day 1 did not significantly differ across groups. (n = 8-9 male mice / group). Error bars represent ± SEM. * $p < 0.05$; **** $p < 0.0001$. Veh, vehicle; TAM, tamoxifen; FST, forced swim test; EPM, elevated plus maze; CFC, contextual fear conditioning; Avg., average; sec, seconds; min, minutes; Sal, saline; K, (*R,S*)-ketamine (30 mg/kg).

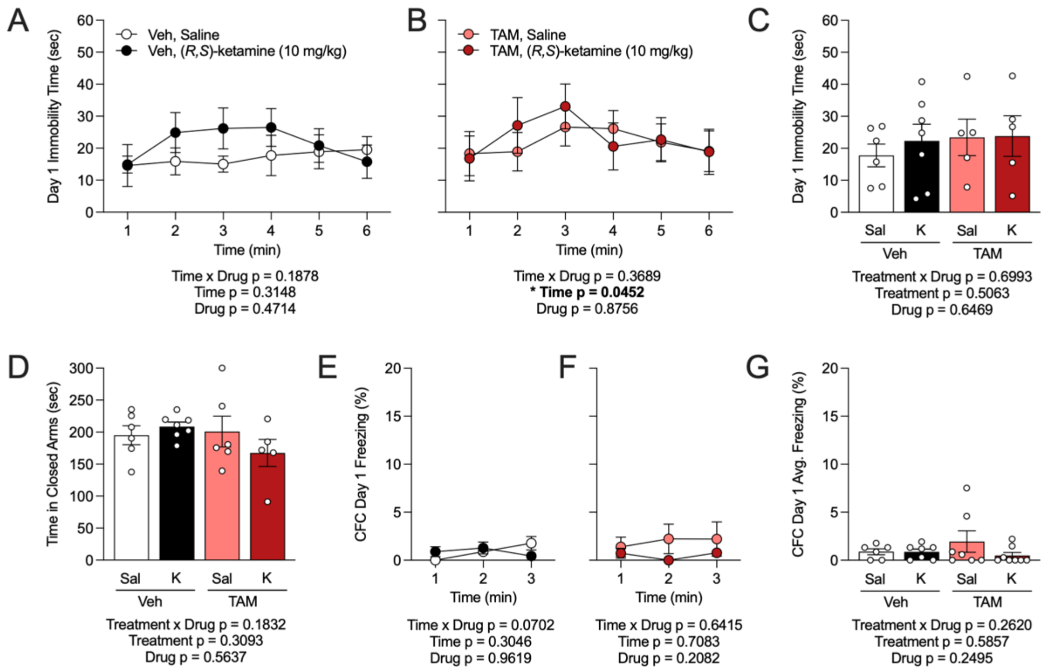


**Supplementary Fig. 2. Deletion of NR2B from 6-week-old adult-born neurons does not alter (*R,S*)-ketamine efficacy in female mice.** (**A**) On day 1 of the FST, there was no difference in immobility time between Veh-Sal and Veh-K mice or (**B**) between TAM-Sal and TAM-K mice. (**C**) Average immobility time in the FST did not differ across groups. (**D**) Time spent in the closed arms of the EPM did not differ across groups. (**E**) Freezing on day 1 of CFC did not differ between Veh-Sal and Veh-K or (**F**) between TAM-Sal and TAM-K mice. (**G**) Average freezing during CFC Day 1 did not significantly differ across groups. (n = 5-8 female mice / group). Error bars represent ± SEM. * $p < 0.05$. Veh, vehicle; TAM, tamoxifen; FST, forced swim test; EPM, elevated plus maze; CFC, contextual fear conditioning; Avg., average; sec, seconds; min, minutes; Sal, saline; K, (*R,S*)-ketamine (10 mg/kg).

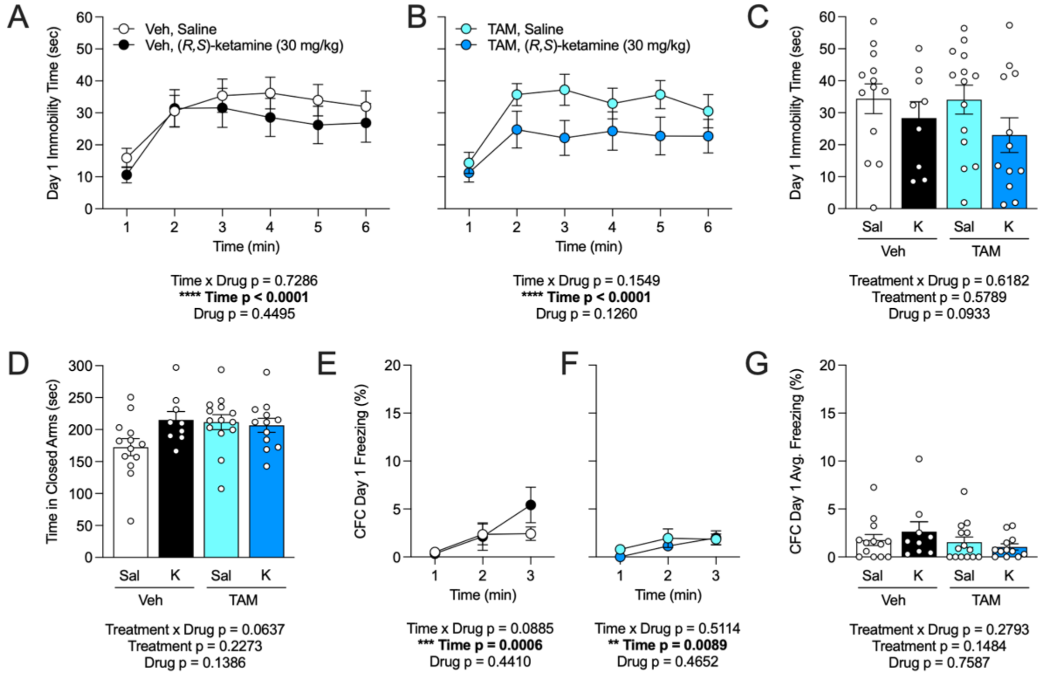


**Supplementary Fig. 3. Deletion of NR2B from 2-week-old adult-born neurons does not alter (*R,S*)-ketamine efficacy in male mice.** (**A**) On day 1 of the FST, there was no difference in immobility time between Veh-Sal and Veh-K mice or (**B**) between TAM-Sal and TAM-K mice. (**C**) Average immobility time in the FST did not differ across groups. (**D**) Time spent in the closed arms of the EPM did not differ between groups. (**E**) Freezing on day 1 of the CFC did not differ between Veh-Sal and Veh-K or (**F**) between TAM-Sal and TAM-K mice. (**G**) Average freezing during day 1 of CFC did not significantly differ across groups. (n = 8-14 male mice / group). Error bars represent ± SEM. ** $p < 0.01$; *** $p < 0.001$; **** $p < 0.0001$. Veh, vehicle; TAM, tamoxifen; FST, forced swim test; EPM, elevated plus maze; NSF, novelty suppressed feeding; CFC, contextual fear conditioning; Avg., average; sec, seconds; min, minutes; g, grams; Sal, saline; K, (*R,S*)-ketamine (30 mg/kg).

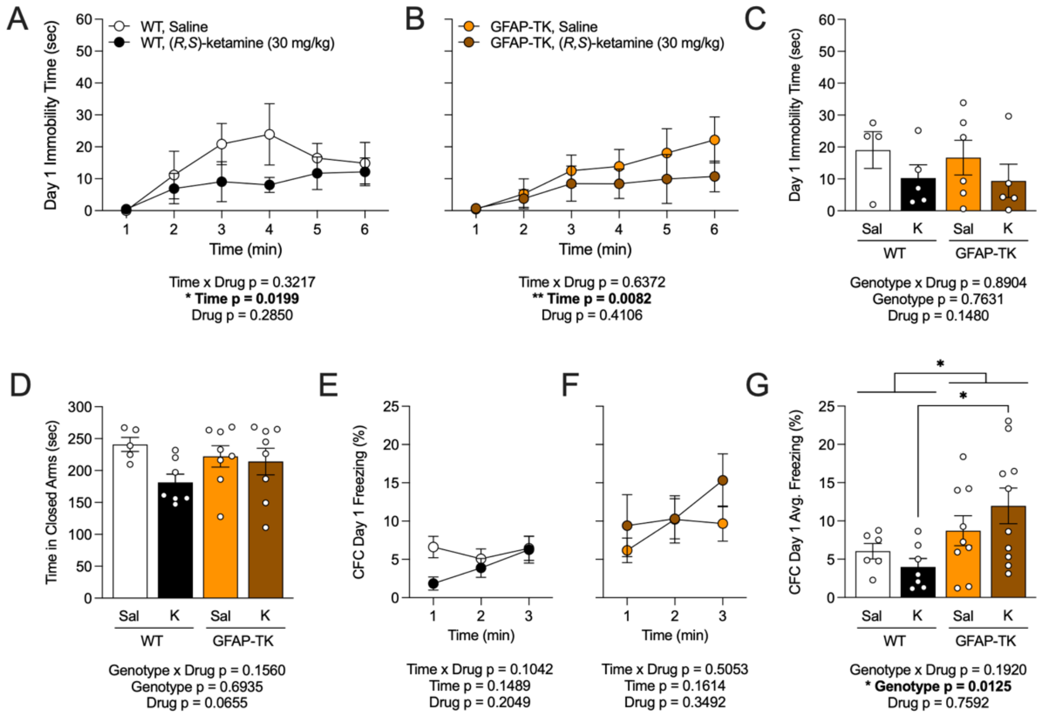


**Supplementary Fig. 4. Ablation of adult neurogenesis does not alter behavioral despair or anxiety-like behavior.** (**A**) On day 1 of the FST, there was no difference in immobility time between WT Sal and WT K mice, or (**B**) between GFAP-TK Sal and GFAP-TK K mice. (**C**) Average immobility time in the FST did not differ across groups. (**D**) Time spent in the closed arms of the EPM did not differ across groups. (**E**) Freezing on day 1 of the CFC did not differ between WT Sal or WT K or (**F**) between GFAP-TK Sal and GFAP-TK K mice. (**G**) There is a main effect of Genotype on average freezing during day 1 of CFC. GFAP-TK K mice froze more than WT K mice. (n = 6-9 male mice / group). Error bars represent ± SEM. * $p < 0.05$; ** $p < 0.01$. WT, wild-type; GFAP-TK, glial fibrillary acidic protein-thymidine kinase; FST, forced swim test; EPM, elevated plus maze; CFC, contextual fear conditioning; Avg., average; sec, seconds; min, minutes; K, (*R,S*)-ketamine (30 mg/kg).

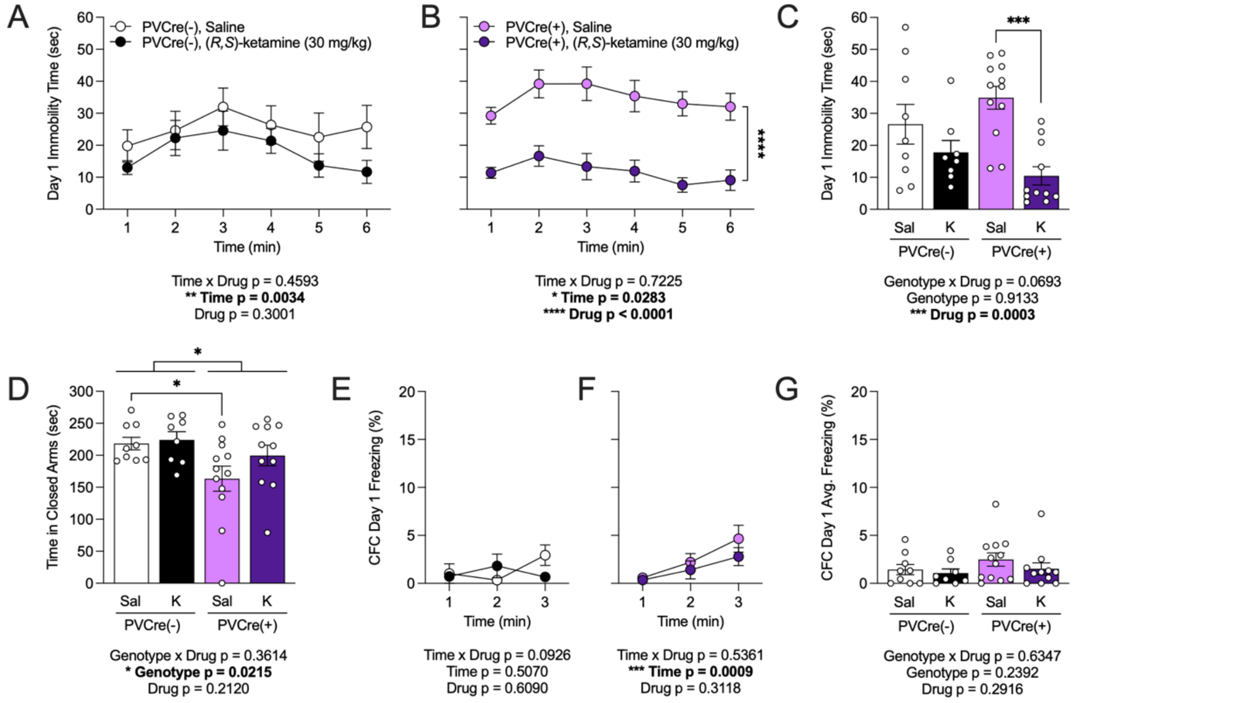


**Supplementary Fig. 5. Deletion of NR2B from inhibitory interneurons selectively alters (*R,S*)-ketamine efficacy in male mice.** (**A**) On day 1 of the FST, there was no difference in immobility time between PV-Cre(-) Sal and PV-Cre(-) K mice. (**B**) However, PV-Cre(+) K mice spent significantly less time immobile than PV-Cre(+) Sal mice. (**C**) K reduced average immobility time in PV-Cre(+) mice but had no effect on PV-Cre(-) mice. (**D**) PV-Cre(+) mice spent less time in the closed arms of the EPM as compared to PV-Cre(-) mice. PV-Cre(+) Sal mice spent less time in the closed arms as compared to PV-Cre(-) Sal mice. (**E**) There was no effect of K on freezing during day 1 of the CFC on PV-Cre(-) or (**F**) PV-Cre(+) mice. (**G**) Average freezing on day 1 of the CFC did not differ across groups. (n = 8-12 male mice / group). Error bars represent ± SEM. * $p < 0.05$; ** $p < 0.01$; *** $p < 0.001$; **** $p < 0.0001$. PV-Cre, parvalbumin-Cre; FST, forced swim test; EPM, elevated plus maze; CFC, contextual fear conditioning; Avg., average; sec, seconds; min, minutes; Sal, saline; K, (*R,S*)-ketamine (30 mg/kg).

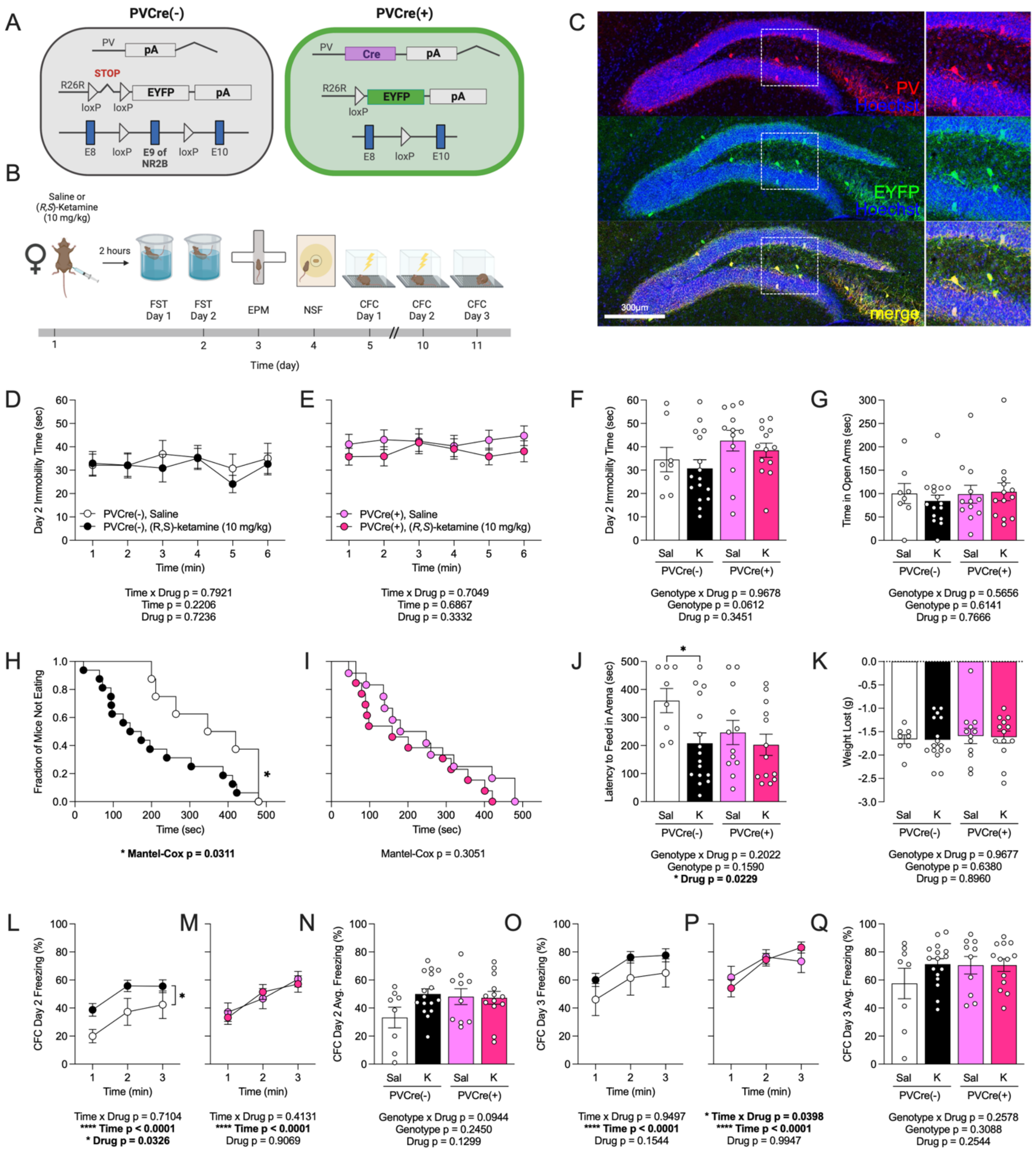

**Supplementary Fig. 6. Deletion of NR2B from inhibitory interneurons selectively influences (*R,S*)-ketamine efficacy in female mice**. (**A**) Genetic schematic. (**B**) Experimental timeline. (**C**) Representative section showing PV (red), EYFP (green), and Hoechst (blue) labeling in the HPC of PV-Cre x NR2B$^{f/f}$ x EYFP female mice, indicating PV-expressing cells with the NR2B subunit deleted. (**D**) In PV-Cre(-) mice and (**E**) PV-Cre(+) mice, there was no effect of K on immobility time during the FST. (**F**) There was no difference in average immobility time in the FST across groups. (**G**) Time in the open arms of the EPM did not differ across groups. (**H**) In the NSF, K-administered PV-Cre(-) mice approached the food quicker than Sal-administered PV-Cre(-) mice. (**I**) There was no effect of K on latency to feed in the NSF in PV-Cre(+) mice. (**J**) K-administered PV-Cre(-) mice had a lower latency to feed than Sal-administered PV-Cre(-) mice, whereas there was no effect of K in PV-Cre(+) mice. (**K**) There was no difference in weight loss in the NSF across groups. (**L**) The percentage of freezing on day 2 of the CFC was increased in K-administered PV-Cre(-) mice as compared to Sal-administered PV-Cre(-) mice. (**M**) PV-Cre(+) Sal- or K-treated mice had comparable levels of freezing on day 2 of the CFC. (**N**) Average freezing on day 2 of the CFC did not differ across groups. (**O**) The percentage of freezing did not differ among PV-Cre(-) or (**P**) PV-Cre(+) Sal- or K-treated mice on day 3 of the CFC, a trend which was reflected in (**Q**) average freezing across groups. (n = 8-16 female mice / group). Error bars represent ± SEM. * $p < 0.05$; **** $p < 0.0001$. PV, parvalbumin; EYFP, enhanced yellow fluorescent protein; HPC, hippocampus; FST, forced swim test; EPM, elevated plus maze; NSF, novelty suppressed feeding; CFC, contextual fear conditioning; sec, seconds; min, minutes; g, grams; K, (*R,S*)-ketamine (10 mg/kg).

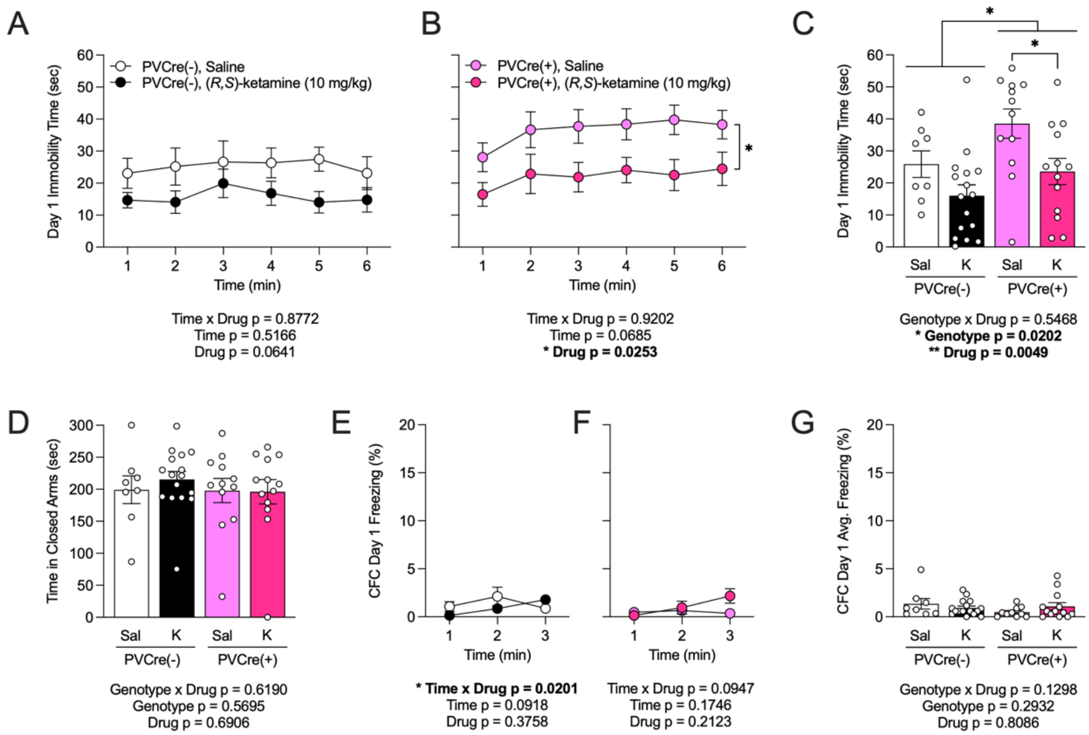


**Supplementary Fig. 7. Deletion of NR2B from inhibitory interneurons increases behavioral despair in female mice, which is alleviated by (*R,S*)-ketamine administration.** (**A**) On day 1 of the FST, in PV-Cre(-) mice, there was no effect of K on immobility time during the FST. (**B**) However, in PV-Cre(+) mice, K mice were spent less time immobile than Sal mice. (**C**) PV-Cre(+) mice spent more average time immobile as compared to PV-Cre(-) mice. K decreased average immobility time in PV-Cre(+) but not PV-Cre(-) mice. (**D**) There was no difference in time spent in the closed arms of the EPM across groups. (**E**) There was no effect of K on freezing during day 1 of the CFC on PV-Cre(-) or (**F**) PV-Cre(+) mice. (**G**) Average freezing on day 1 of the CFC did not differ across groups. (n = 8-12 male mice / group). Error bars represent ± SEM. * $p < 0.05$; ** $p < 0.01$. PV-Cre, parvalbumin-Cre; FST, forced swim test; EPM, elevated plus maze; CFC, contextual fear conditioning; Avg., average; sec, seconds; min, minutes; g, grams; Sal, saline; K, (*R,S*)-ketamine (10 mg/kg).

**Supplemental Table 1. Statistical analyses.**

**Supplemental Table 2. Key resources.**
