## Supplemental Table 1 for "(*R,S*)-ketamine’s rapid-acting antidepressant effects are modulated by NR2B-containing NMDA receptors on adult-born hippocampal neurons"

| Genetic Line / Cohort | Strain | Sex | Behavioral Paradigm | Measurement | Statistical Test | Group | Comparison | F, t, or χ2 | p | * | Fig. |
|---|---|---|---|---|---|---|---|---|---|---|---|
| NestinCreERT2 (6 week) | C57BL/6J x 129S6/SvEv (mixed) | Male (M) | Forced Swim Test (FST) Day 1 | Immobility time per min (sec) | 3way RMANOVA | - | **Time** | **F (5, 165) = 4.826** | **0.0004** | ******* | S1A-B |
| | | | | | | | Treatment | F (1, 33) = 0.1185 | 0.7329 | ns | |
| | | | | | | | **Drug** | **F (1, 33) = 5.203** | **0.0291** | ***** | |
| | | | | | | | Time x Treatment | F (5, 165) = 0.4169 | 0.8365 | ns | |
| | | | | | | | **Time x Drug** | **F (5, 165) = 3.254** | **0.0079** | ****** | |
| | | | | | | | Treatment x Drug | F (1, 33) = 0.07631 | 0.7841 | ns | |
| | | | | | | | Time x Treatment x Drug | F (5, 165) = 0.2443 | 0.9422 | ns | |
| | | | | | 2way RMANOVA | Vehicle | Time x Drug | F (5, 75) = 1.398 | 0.2348 | ns | S1A |
| | | | | | | | Time | F (2.381, 35.71) = 1.814 | 0.1719 | ns | |
| | | | | | | | Drug | F (1, 15) = 2.925 | 0.1078 | ns | |
| | | | | | | Tamoxifen | Time x Drug | F (5, 90) = 2.238 | 0.0572 | ns | S1B |
| | | | | | | | **Time** | **F (2.209, 39.76) = 3.966** | **0.0235** | ***** | |
| | | | | | | | Drug | F (1, 18) = 2.249 | 0.1511 | ns | |
| | | | | Average immobility time (sec) | 2way ANOVA | - | Treatment x Drug | F (1, 33) = 0.02837 | 0.8673 | ns | S1C |
| | | | | | | | Treatment | F (1, 33) = 0.1336 | 0.7171 | ns | |
| | | | | | | | **Drug** | **F (1, 33) = 5.069** | **0.0311** | ***** | |
| | | | | | Šidák's | Vehicle | Drug | t = 1.648 | 0.2058 | ns | |
| | | | | | | Tamoxifen | Drug | t = 1.534 | 0.2511 | ns | |
| | | | FST Day 2 | Immobility time per min (sec) | 3way RMANOVA | - | **Time** | **F (3.229, 113.0) = 14.47** | **<0.0001** | ******** | 1D-E |
| | | | | | | | Treatment | F (1, 35) = 0.007475 | 0.9316 | ns | |
| | | | | | | | **Drug** | **F (1, 35) = 14.24** | **0.0006** | ******* | |
| | | | | | | | Time x Treatment | F (5, 175) = 0.3869 | 0.8574 | ns | |
| | | | | | | | Time x Drug | F (5, 175) = 0.2902 | 0.9179 | ns | |
| | | | | | | | Treatment x Drug | F (1, 35) = 1.175 | 0.2858 | ns | |
| | | | | | | | Time x Treatment x Drug | F (5, 175) = 0.2001 | 0.9621 | ns | |
| | | | | | 2way RMANOVA | Vehicle | Time x Drug | F (5, 80) = 0.2587 | 0.9342 | ns | 1D |
| | | | | | | | **Time** | **F (2.642, 42.27) = 5.563** | **0.0037** | ****** | |
| | | | | | | | **Drug** | **F (1, 16) = 12.17** | **0.0030** | ****** | |
| | | | | | | Tamoxifen | Time x Drug | F (5, 95) = 0.2059 | 0.9593 | ns | 1E |
| | | | | | | | **Time** | **F (2.781, 52.84) = 9.948** | **<0.0001** | ******** | |
| | | | | | | | Drug | F (1, 19) = 3.608 | 0.0728 | ns | |
| | | | | Average immobility time (sec) | 2way ANOVA | - | Treatment x Drug | F (1, 35) = 0.6866 | 0.4129 | ns | 1F |
| | | | | | | | Treatment | F (1, 35) = 0.0006711 | 0.9795 | ns | |
| | | | | | | | **Drug** | **F (1, 35) = 9.166** | **0.0046** | ****** | |
| | | | | | Šidák's | Vehicle | **Drug** | **t = 2.633** | **0.0249** | ***** | |
| | | | | | | Tamoxifen | Drug | t = 1.615 | 0.2174 | ns | |
| | | | Elevated Plus Maze (EPM) | Time in open arms (sec) | 2way ANOVA | - | Treatment x Drug | F (1, 37) = 0.7949 | 0.3784 | ns | 1G |
| | | | | | | | Treatment | F (1, 37) = 0.4045 | 0.5287 | ns | |
| | | | | | | | Drug | F (1, 37) = 1.301 | 0.2613 | ns | |
| | | | | Time in closed arms (sec) | 2way ANOVA | - | Treatment x Drug | F (1, 37) = 0.8914 | 0.3512 | ns | S1D |
| | | | | | | | Treatment | F (1, 37) = 0.9126 | 0.3456 | ns | |
| | | | | | | | Drug | F (1, 37) = 1.114 | 0.2980 | ns | |
| | | | Novelty Suppressed Feeding (NSF) | Fraction of mice not eating | Kaplan-Meier | - | **Treatment x Drug** | **χ2 = 35.28** | **<0.0001** | ******** | 1H-I |
| | | | | | | Vehicle | **Drug** | **χ2 = 8.875** | **0.0029** | ****** | 1H |
| | | | | | | Tamoxifen | Drug | χ2 = 0.7173 | 0.3970 | ns | 1I |
| | | | | Latency to feed in arena (sec) | 2way ANOVA | - | **Treatment x Drug** | **F (1, 33) = 5.832** | **0.0214** | ***** | 1J |
| | | | | | | | **Treatment** | **F (1, 33) = 18.85** | **0.0001** | ******* | |
| | | | | | | | Drug | F (1, 33) = 3.440 | 0.0726 | ns | |
| | | | | | Šidák's | Vehicle | **Drug** | **t = 2.837** | **0.0455** | ***** | |
| | | | | | | Saline | Treatment | t = 1.463 | 0.6304 | ns | |
| | | | | | | - | Veh:Sal vs. TAM:K30 | t = 1.767 | 0.4190 | ns | |
| | | | | | | | **Veh:K30 vs. TAM:Sal** | **t = 4.363** | **0.0007** | ******* | |
| | | | | | | (*R*,*S*)-ketamine | **Treatment** | **t = 4.489** | **0.0005** | ******* | |
| | | | | | | Tamoxifen | Drug | t = 0.4254 | 0.9988 | ns | |
| | | | | Weight lost (g) | 2way ANOVA | - | Treatment x Drug | F (1, 34) = 0.7468 | 0.3936 | ns | 1K |
| | | | | | | | Treatment | F (1, 34) = 0.2359 | 0.6303 | ns | |
| | | | | | | | Drug | F (1, 34) = 0.6150 | 0.4384 | ns | |
| | | | Contextual Fear Conditioning (CFC) Day 1 | Freezing per min (%) | 3way RMANOVA | - | **Time** | **F (1.691, 57.50) = 15.25** | **<0.0001** | ******** | S1E-F |
| | | | | | | | Treatment | F (1, 34) = 0.08625 | 0.7708 | ns | |
| | | | | | | | Drug | F (1, 34) = 2.520 | 0.1217 | ns | |
| | | | | | | | Time x Treatment | F (2, 68) = 1.752 | 0.1812 | ns | |
| | | | | | | | Time x Drug | F (2, 68) = 1.017 | 0.3670 | ns | |
| | | | | | | | Treatment x Drug | F (1, 34) = 2.608 | 0.1156 | ns | |
| | | | | | | | Time x Treatment x Drug | F (2, 68) = 0.07864 | 0.9245 | ns | |
| | | | | | 2way RMANOVA | Vehicle | Time x Drug | F (2, 30) = 0.6658 | 0.5213 | ns | S1E |
| | | | | | | | Time | F (1.835, 27.53) = 2.770 | 0.0842 | ns | |
| | | | | | | | Drug | F (1, 15) = 3.362 | 0.0866 | ns | |
| | | | | | | Tamoxifen | Time x Drug | F (2, 38) = 0.3443 | 0.7109 | ns | S1F |
| | | | | | | | **Time** | **F (1.437, 27.29) = 16.56** | **<0.0001** | ******** | |
| | | | | | | | Drug | F (1, 19) = 0.0005976 | 0.9808 | ns | |
| | | | | Average freezing (%) | 2way ANOVA | - | Treatment x Drug | F (1, 34) = 3.419 | 0.0732 | ns | S1G |
| | | | | | | | Treatment | F (1, 34) = 0.004931 | 0.9444 | ns | |
| | | | | | | | Drug | F (1, 34) = 1.881 | 0.1793 | ns | |
| | | | CFC Day 2 | Freezing per min (%) | 3way RMANOVA | - | **Time** | **F (1.742, 59.24) = 4.530** | **0.0186** | ***** | 1L-M |
| | | | | | | | Treatment | F (1, 34) = 1.713 | 0.1994 | ns | |
| | | | | | | | Drug | F (1, 34) = 2.502 | 0.1229 | ns | |
| | | | | | | | Time x Treatment | F (2, 68) = 0.5038 | 0.6064 | ns | |
| | | | | | | | Time x Drug | F (2, 68) = 1.741 | 0.1831 | ns | |
| | | | | | | | Treatment x Drug | F (1, 34) = 3.081 | 0.0882 | ns | |
| | | | | | | | Time x Treatment x Drug | F (2, 68) = 2.280 | 0.1101 | ns | |
| | | | | | 2way RMANOVA | Vehicle | Time x Drug | F (2, 30) = 3.278 | 0.0516 | ns | 1L |
| | | | | | | | Time | F (1.479, 22.18) = 0.9729 | 0.3692 | ns | |
| | | | | | | | Drug | F (1, 15) = 4.100 | 0.0611 | ns | |
| | | | | | | Tamoxifen | Time x Drug | F (2, 38) = 0.07323 | 0.9295 | ns | 1M |
| | | | | | | | **Time** | **F (1.931, 36.69) = 4.644** | **0.0168** | ***** | |
| | | | | | | | Drug | F (1, 19) = 0.02051 | 0.8876 | ns | |
| | | | | Average freezing (%) | 2way ANOVA | - | Treatment x Drug | F (1, 34) = 3.081 | 0.0882 | ns | 1N |
| | | | | | | | Treatment | F (1, 34) = 1.713 | 0.1994 | ns | |
| | | | | | | | Drug | F (1, 34) = 2.502 | 0.1229 | ns | |
| | | | CFC Day 3 | Freezing per min (%) | 3way RMANOVA | - | **Time** | **F (1.684, 57.24) = 18.90** | **<0.0001** | ******** | 1O-P |
| | | | | | | | Treatment | F (1, 34) = 0.001755 | 0.9668 | ns | |
| | | | | | | | Drug | F (1, 34) = 0.7071 | 0.4063 | ns | |
| | | | | | | | Time x Treatment | F (2, 68) = 1.320 | 0.2738 | ns | |
| | | | | | | | Time x Drug | F (2, 68) = 0.2826 | 0.7547 | ns | |
| | | | | | | | **Treatment x Drug** | **F (1, 34) = 5.321** | **0.0273** | ***** | |
| | | | | | | | Time x Treatment x Drug | F (2, 68) = 0.3342 | 0.7171 | ns | |
| | | | | | 2way RMANOVA | Vehicle | Time x Drug | F (2, 30) = 0.001268 | 0.9987 | ns | 1O |
| | | | | | | | **Time** | **F (1.464, 21.97) = 5.237** | **0.0212** | ***** | |
| | | | | | | | **Drug** | **F (1, 15) = 5.628** | **0.0315** | ***** | |
| | | | | | | Tamoxifen | Time x Drug | F (2, 38) = 0.6558 | 0.5248 | ns | 1P |
| | | | | | | | **Time** | **F (1.803, 34.27) = 15.73** | **<0.0001** | ******** | |
| | | | | | | | Drug | F (1, 19) = 1.029 | 0.3231 | ns | |
| | | | | Average freezing | 2way ANOVA | - | **Treatment x Drug** | **F (1, 33) = 7.806** | **0.0086** | ****** | |
| | | | | | | | Treatment | F (1, 33) = 0.1431 | 0.7076 | ns | |

| | | | | | | | | | | | |
|---|---|---|---|---|---|---|---|---|---|---|---|
| | | | | Average freezing (%) | | | Drug | F (1, 33) = 1.655 | 0.2073 | ns | 1Q |
| | | | | | Šidák's | Vehicle | **Drug** | **t = 2.711** | **0.0210** | ***** | |
| | | | | | | Tamoxifen | Drug | t = 1.145 | 0.5075 | ns | |

| Genetic Line / Cohort | Strain | Sex | Behavioral Paradigm | Measurement | Statistical Test | Group | Comparison | F, t, or $\chi^2$ | p | * | Fig. |
|---|---|---|---|---|---|---|---|---|---|---|---|
| NestinCreERT2 (6 week) | 129S6/SvEv | Female (F) | FST Day 1 | Immobility time per min (sec) | 3way RMANOVA | - | **Time** | **F (5, 95) = 3.510** | **0.0059** | ****** | S2A-B |
| | | | | | | | Treatment | F (1, 19) = 0.4168 | 0.5262 | ns | |
| | | | | | | | Drug | F (1, 19) = 0.3403 | 0.5665 | ns | |
| | | | | | | | Time x Treatment | F (5, 95) = 0.7528 | 0.5861 | ns | |
| | | | | | | | Time x Drug | F (5, 95) = 1.805 | 0.1192 | ns | |
| | | | | | | | Treatment x Drug | F (1, 19) = 0.09830 | 0.7573 | ns | |
| | | | | | | | Time x Treatment x Drug | F (5, 95) = 0.7750 | 0.5701 | ns | |
| | | | | | 2way RMANOVA | Vehicle | Time x Drug | F (5, 55) = 1.556 | 0.1878 | ns | S2A |
| | | | | | | | Time | F (3.438, 37.82) = 1.227 | 0.3148 | ns | |
| | | | | | | | Drug | F (1, 11) = 0.5562 | 0.4714 | ns | |
| | | | | | | Tamoxifen | Time x Drug | F (5, 40) = 1.113 | 0.3689 | ns | S2B |
| | | | | | | | **Time** | **F (2.952, 23.61) = 3.133** | **0.0452** | ***** | |
| | | | | | | | Drug | F (1, 8) = 0.02614 | 0.8756 | ns | |
| | | | | Average immobility time (sec) | 2way ANOVA | - | Treatment x Drug | F (1, 19) = 0.1538 | 0.6993 | ns | S2C |
| | | | | | | | Treatment | F (1, 19) = 0.4588 | 0.5063 | ns | |
| | | | | | | | Drug | F (1, 19) = 0.2167 | 0.6469 | ns | |
| | | | FST Day 2 | Immobility time per min (sec) | 3way RMANOVA (mixed-effects anaysis) | - | **Time** | **F (3.788, 69.71) = 4.558** | **0.0030** | ****** | 2D-E |
| | | | | | | | Treatment | F (1, 20) = 1.353 | 0.2584 | ns | |
| | | | | | | | Drug | F (1, 20) = 0.2825 | 0.6009 | ns | |
| | | | | | | | **Time x Treatment** | **F (5, 92) = 4.143** | **0.0019** | ****** | |
| | | | | | | | Time x Drug | F (5, 92) = 1.773 | 0.1260 | ns | |
| | | | | | | | Treatment x Drug | F (1, 20) = 0.3485 | 0.5616 | ns | |
| | | | | | | | Time x Treatment x Drug | F (5, 92) = 0.3826 | 0.8596 | ns | |
| | | | | | 2way RMANOVA (mixed-effects analysis) | Vehicle | Time x Drug | F (5, 47) = 1.144 | 0.3509 | ns | 2D |
| | | | | | | | **Time** | **F (2.973, 27.95) = 6.972** | **0.0012** | ****** | |
| | | | | | | | Drug | F (1, 11) = 0.6944 | 0.4224 | ns | |
| | | | | | 2way RMANOVA | Tamoxifen | Time x Drug | F (5, 45) = 1.058 | 0.3961 | ns | 2E |
| | | | | | | | Time | F (2.872, 25.85) = 0.9045 | 0.4488 | ns | |
| | | | | | | | Drug | F (1, 9) = 0.001600 | 0.9690 | ns | |
| | | | | Average immobility time (sec) | 2way ANOVA | - | Treatment x Drug | F (1, 20) = 0.4067 | 0.5309 | ns | 2F |
| | | | | | | | Treatment | F (1, 20) = 0.2533 | 0.6203 | ns | |
| | | | | | | | Drug | F (1, 20) = 0.08482 | 0.7739 | ns | |
| | | | EPM | Time in open arms (sec) | 2way ANOVA | - | Treatment x Drug | F (1, 20) = 1.867 | 0.1869 | ns | 2G |
| | | | | | | | Treatment | F (1, 20) = 1.118 | 0.3029 | ns | |
| | | | | | | | Drug | F (1, 20) = 0.3385 | 0.5672 | ns | |
| | | | | Time in closed arms (sec) | 2way ANOVA | - | Treatment x Drug | F (1, 20) = 1.901 | 0.1832 | ns | S2D |
| | | | | | | | Treatment | F (1, 20) = 1.088 | 0.3093 | ns | |
| | | | | | | | Drug | F (1, 20) = 0.3447 | 0.5637 | ns | |
| | | | NSF | Fraction of mice not eating | Kaplan-Meier | - | **Treatment x Drug** | **$\chi^2$ = 10.28** | **0.0163** | ***** | 2H-I |
| | | | | | | Vehicle | **Drug** | **$\chi^2$ = 5.217** | **0.0224** | ***** | 2H |
| | | | | | | Tamoxifen | Drug | $\chi^2$ = 3.114 | 0.0776 | ns | 2I |
| | | | | Latency to feed in arena (sec) | 2way ANOVA | - | Treatment x Drug | F (1, 23) = 1.319 | 0.2626 | ns | 2J |
| | | | | | | | Treatment | F (1, 23) = 0.2106 | 0.6506 | ns | |
| | | | | | | | **Drug** | **F (1, 23) = 9.113** | **0.0061** | ****** | |
| | | | | | Šidák's | Vehicle | **Drug** | **t = 2.781** | **0.0211** | ***** | |
| | | | | | | Tamoxifen | Drug | t = 1.412 | 0.3133 | ns | |
| | | | | Weight lost (g) | 2way ANOVA | - | Treatment x Drug | F (1, 23) = 0.6425 | 0.4310 | ns | 2K |
| | | | | | | | Treatment | F (1, 23) = 0.02001 | 0.8887 | ns | |
| | | | | | | | **Drug** | **F (1, 23) = 6.245** | **0.0200** | ***** | |
| | | | | | Šidák's | Vehicle | Drug | t = 2.203 | 0.0744 | ns | |
| | | | | | | Tamoxifen | Drug | t = 1.282 | 0.3803 | ns | |
| | | | CFC Day 1 | Freezing per min (%) | 3way RMANOVA | - | Time | F (1.323, 31.76) = 0.6423 | 0.4708 | ns | S2E-F |
| | | | | | | | Treatment | F (1, 24) = 0.3064 | 0.5850 | ns | |
| | | | | | | | Drug | F (1, 24) = 1.394 | 0.2492 | ns | |
| | | | | | | | Time x Treatment | F (2, 48) = 0.1745 | 0.8404 | ns | |
| | | | | | | | Time x Drug | F (2, 48) = 1.202 | 0.3094 | ns | |
| | | | | | | | Treatment x Drug | F (1, 24) = 1.317 | 0.2624 | ns | |
| | | | | | | | Time x Treatment x Drug | F (2, 48) = 0.8102 | 0.4508 | ns | |
| | | | | | 2way RMANOVA | Vehicle | Time x Drug | F (2, 22) = 3.005 | 0.0702 | ns | S2E |
| | | | | | | | Time | F (1.979, 21.77) = 1.255 | 0.3046 | ns | |
| | | | | | | | Drug | F (1, 11) = 0.002392 | 0.9619 | ns | |
| | | | | | | Tamoxifen | Time x Drug | F (2, 26) = 0.4515 | 0.6415 | ns | S2F |
| | | | | | | | Time | F (1.098, 14.27) = 0.1714 | 0.7083 | ns | |
| | | | | | | | Drug | F (1, 13) = 1.754 | 0.2082 | ns | |
| | | | | Average freezing (%) | 2way ANOVA | - | Treatment x Drug | F (1, 24) = 1.319 | 0.2620 | ns | S2G |
| | | | | | | | Treatment | F (1, 24) = 0.3053 | 0.5857 | ns | |
| | | | | | | | Drug | F (1, 24) = 1.393 | 0.2495 | ns | |
| | | | CFC Day 2 | Freezing per min (%) | 3way RMANOVA | - | **Time** | **F (1.934, 46.41) = 12.51** | **<0.0001** | ******** | 2L-M |
| | | | | | | | Treatment | F (1, 24) = 2.393 | 0.1350 | ns | |
| | | | | | | | Drug | F (1, 24) = 0.7421 | 0.3975 | ns | |
| | | | | | | | Time x Treatment | F (2, 48) = 0.1147 | 0.8919 | ns | |
| | | | | | | | Time x Drug | F (2, 48) = 0.4253 | 0.6560 | ns | |
| | | | | | | | Treatment x Drug | F (1, 24) = 0.2377 | 0.6303 | ns | |
| | | | | | | | Time x Treatment x Drug | F (2, 48) = 1.932 | 0.1560 | ns | |
| | | | | | 2way RMANOVA | Vehicle | Time x Drug | F (2, 22) = 1.925 | 0.1696 | ns | 2L |
| | | | | | | | **Time** | **F (1.804, 19.85) = 6.241** | **0.0094** | ****** | |
| | | | | | | | Drug | F (1, 11) = 1.696 | 0.2195 | ns | |
| | | | | | | Tamoxifen | Time x Drug | F (2, 26) = 0.3058 | 0.7391 | ns | 2M |
| | | | | | | | **Time** | **F (1.911, 24.84) = 6.388** | **0.0063** | ****** | |
| | | | | | | | Drug | F (1, 13) = 0.05294 | 0.8216 | ns | |
| | | | | Average freezing (%) | 2way ANOVA | - | Treatment x Drug | F (1, 24) = 0.2379 | 0.6302 | ns | 2N |
| | | | | | | | Treatment | F (1, 24) = 2.393 | 0.1350 | ns | |
| | | | | | | | Drug | F (1, 24) = 0.7417 | 0.3976 | ns | |
| | | | CFC Day 3 | Freezing per min (%) | 3way RMANOVA | - | **Time** | **F (1.896, 45.50) = 25.86** | **<0.0001** | ******** | 2O-P |
| | | | | | | | Treatment | F (1, 24) = 2.188 | 0.1521 | ns | |
| | | | | | | | **Drug** | **F (1, 24) = 4.741** | **0.0395** | ***** | |
| | | | | | | | **Time x Treatment** | **F (2, 48) = 4.178** | **0.0212** | ***** | |
| | | | | | | | Time x Drug | F (2, 48) = 0.6517 | 0.5257 | ns | |
| | | | | | | | Treatment x Drug | F (1, 24) = 0.5822 | 0.4529 | ns | |
| | | | | | | | Time x Treatment x Drug | F (2, 48) = 1.353 | 0.2682 | ns | |
| | | | | | 2way RMANOVA | Vehicle | Time x Drug | F (2, 22) = 1.502 | 0.2447 | ns | 2O |
| | | | | | | | **Time** | **F (1.843, 20.28) = 4.741** | **0.0226** | ***** | |
| | | | | | | | Drug | F (1, 11) = 3.665 | 0.0819 | ns | |
| | | | | | | Tamoxifen | Time x Drug | F (2, 26) = 0.3994 | 0.6748 | ns | 2P |
| | | | | | | | **Time** | **F (1.793, 23.31) = 27.21** | **<0.0001** | ******** | |
| | | | | | | | Drug | F (1, 13) = 1.178 | 0.2975 | ns | |
| | | | | Average freezing (%) | 2way ANOVA | - | Treatment x Drug | F (1, 24) = 0.5819 | 0.4530 | ns | 2Q |
| | | | | | | | Treatment | F (1, 24) = 2.188 | 0.1521 | ns | |
| | | | | | | | **Drug** | **F (1, 24) = 4.741** | **0.0395** | ***** | |
| | | | | | Šidák's | Vehicle | Drug | t = 2.008 | 0.1090 | ns | |

| | | | | | | | | | | |
|---|---|---|---|---|---|---|---|---|---|---|
| | | | | | Šidák's | Tamoxifen | Drug | t = 1.038 | 0.5231 | ns |

| Genetic Line / Cohort | Strain | Sex | Behavioral Paradigm | Measurement | Statistical Test | Group | Comparison | F, t, or χ2 | p | * | Fig. |
|---|---|---|---|---|---|---|---|---|---|---|---|
| NestinCreERT2 (2 week) | C57BL/6J x 129S6/SvEv (mixed) | M | FST Day 1 | Immobility time per min (sec) | 3way RMANOVA | - | **Time** | **F (5, 220) = 28.43** | **<0.0001** | ******** | S3A-B |
| | | | | | | | Treatment | F (1, 44) = 0.2217 | 0.6401 | ns | |
| | | | | | | | Drug | F (1, 44) = 2.723 | 0.1060 | ns | |
| | | | | | | | Time x Treatment | F (5, 220) = 0.3471 | 0.8838 | ns | |
| | | | | | | | Time x Drug | F (5, 220) = 0.8592 | 0.5094 | ns | |
| | | | | | | | Treatment x Drug | F (1, 44) = 0.3170 | 0.5763 | ns | |
| | | | | | | | Time x Treatment x Drug | F (5, 220) = 1.137 | 0.3416 | ns | |
| | | | | | 2way RMANOVA | Vehicle | Time x Drug | F (5, 100) = 0.5624 | 0.7286 | ns | S3A |
| | | | | | | | **Time** | **F (3.688, 73.75) = 12.57** | **<0.0001** | ******** | |
| | | | | | | | Drug | F (1, 20) = 0.5951 | 0.4495 | ns | |
| | | | | | | Tamoxifen | Time x Drug | F (5, 120) = 1.639 | 0.1549 | ns | S3B |
| | | | | | | | **Time** | **F (3.338, 80.11) = 16.32** | **<0.0001** | ******** | |
| | | | | | | | Drug | F (1, 24) = 2.513 | 0.1260 | ns | |
| | | | | Average immobility time (sec) | 2way ANOVA | - | Treatment x Drug | F (1, 44) = 0.2519 | 0.6182 | ns | S3C |
| | | | | | | | Treatment | F (1, 44) = 0.3127 | 0.5789 | ns | |
| | | | | | | | Drug | F (1, 44) = 2.942 | 0.0933 | ns | |
| | | | FST Day 2 | Immobility time per min (sec) | 3way RMANOVA (mixed-effects anaysis) | - | **Time** | **F (3.792, 163.8) = 38.48** | **<0.0001** | ******** | 3D-E |
| | | | | | | | Treatment | F (1, 44) = 0.2532 | 0.6174 | ns | |
| | | | | | | | Drug | F (1, 44) = 0.6057 | 0.4406 | ns | |
| | | | | | | | Time x Treatment | F (5, 216) = 1.828 | 0.1085 | ns | |
| | | | | | | | Time x Drug | F (5, 216) = 1.918 | 0.0925 | ns | |
| | | | | | | | Treatment x Drug | F (1, 44) = 0.2191 | 0.6421 | ns | |
| | | | | | | | Time x Treatment x Drug | F (5, 216) = 1.684 | 0.1397 | ns | |
| | | | | | 2way RMANOVA (mixed-effects analysis) | Vehicle | Time x Drug | F (5, 96) = 1.825 | 0.1151 | ns | 3D |
| | | | | | | | **Time** | **F (3.433, 65.92) = 11.20** | **<0.0001** | ******** | |
| | | | | | | | Drug | F (1, 20) = 0.03427 | 0.8550 | ns | |
| | | | | | 2way RMANOVA | Tamoxifen | Time x Drug | F (5, 120) = 1.694 | 0.1411 | ns | 3E |
| | | | | | | | **Time** | **F (3.610, 86.65) = 32.32** | **<0.0001** | ******** | |
| | | | | | | | Drug | F (1, 24) = 1.273 | 0.2704 | ns | |
| | | | | Average immobility time (sec) | 2way ANOVA | - | Treatment x Drug | F (1, 44) = 0.0004271 | 0.9836 | ns | 3F |
| | | | | | | | Treatment | F (1, 44) = 0.4657 | 0.4985 | ns | |
| | | | | | | | Drug | F (1, 44) = 0.1668 | 0.6849 | ns | |
| | | | EPM | Time in open arms (sec) | 2way ANOVA | - | Treatment x Drug | F (1, 44) = 4.008 | 0.0515 | ns | 3G |
| | | | | | | | Treatment | F (1, 44) = 1.252 | 0.2693 | ns | |
| | | | | | | | Drug | F (1, 44) = 1.940 | 0.1707 | ns | |
| | | | | Time in closed arms (sec) | 2way ANOVA | - | Treatment x Drug | F (1, 44) = 3.617 | 0.0637 | ns | S3D |
| | | | | | | | Treatment | F (1, 44) = 1.499 | 0.2273 | ns | |
| | | | | | | | Drug | F (1, 44) = 2.275 | 0.1386 | ns | |
| | | | NSF | Fraction of mice not eating | Kaplan-Meier | - | Treatment x Drug | χ2 = 7.529 | 0.0568 | ns | 3H-I |
| | | | | | | Vehicle | Drug | χ2 = 0.005472 | 0.9410 | ns | 3H |
| | | | | | | Tamoxifen | Drug | χ2 = 0.333 | 0.5639 | ns | 3I |
| | | | | Latency to feed in arena (sec) | 2way ANOVA | - | Treatment x Drug | F (1, 44) = 0.002731 | 0.9586 | ns | 3J |
| | | | | | | | **Treatment** | **F (1, 44) = 12.24** | **0.0011** | ****** | |
| | | | | | | | Drug | F (1, 44) = 0.03076 | 0.8616 | ns | |
| | | | | | Šidák's | Saline | **Treatment** | **t = 2.699** | **0.0196** | ***** | |
| | | | | | | (*R,S*)-ketamine | Treatment | t = 2.288 | 0.0533 | ns | |
| | | | | Weight lost (g) | 2way ANOVA | - | Treatment x Drug | F (1, 44) = 0.06685 | 0.7972 | ns | 3K |
| | | | | | | | **Treatment** | **F (1, 44) = 5.863** | **0.0196** | ***** | |
| | | | | | | | Drug | F (1, 44) = 0.004158 | 0.9489 | ns | |
| | | | | | Šidák's | Saline | Treatment | 1.644 | 0.2031 | ns | |
| | | | | | | (*R,S*)-ketamine | Treatment | 1.779 | 0.1575 | ns | |
| | | | CFC Day 1 | Freezing per min (%) | 3way RMANOVA | - | **Time** | **F (1.956, 86.06) = 15.37** | **<0.0001** | ******** | S3E-F |
| | | | | | | | Treatment | F (1, 44) = 2.164 | 0.1483 | ns | |
| | | | | | | | Drug | F (1, 44) = 0.09554 | 0.7587 | ns | |
| | | | | | | | Time x Treatment | F (2, 88) = 2.452 | 0.0920 | ns | |
| | | | | | | | **Time x Drug** | **F (2, 88) = 3.511** | **0.0341** | ***** | |
| | | | | | | | Treatment x Drug | F (1, 44) = 1.200 | 0.2794 | ns | |
| | | | | | | | Time x Treatment x Drug | F (2, 88) = 1.017 | 0.3657 | ns | |
| | | | | | 2way RMANOVA | Vehicle | Time x Drug | F (2, 40) = 2.578 | 0.0885 | ns | S3E |
| | | | | | | | **Time** | **F (1.965, 39.31) = 9.170** | **0.0006** | ******* | |
| | | | | | | | Drug | F (1, 20) = 0.6180 | 0.4410 | ns | |
| | | | | | | Tamoxifen | Time x Drug | F (2, 48) = 0.6802 | 0.5114 | ns | S3F |
| | | | | | | | **Time** | **F (1.813, 43.51) = 5.537** | **0.0089** | ****** | |
| | | | | | | | Drug | F (1, 24) = 0.5508 | 0.4652 | ns | |
| | | | | Average freezing (%) | 2way ANOVA | - | Treatment x Drug | F (1, 44) = 1.200 | 0.2793 | ns | S3G |
| | | | | | | | Treatment | F (1, 44) = 2.164 | 0.1484 | ns | |
| | | | | | | | Drug | F (1, 44) = 0.09554 | 0.7587 | ns | |
| | | | CFC Day 2 | Freezing per min (%) | 3way RMANOVA | - | **Time** | **F (1.845, 81.18) = 21.61** | **<0.0001** | ******** | 3L-M |
| | | | | | | | Treatment | F (1, 44) = 0.2584 | 0.6138 | ns | |
| | | | | | | | Drug | F (1, 44) = 0.06717 | 0.7967 | ns | |
| | | | | | | | Time x Treatment | F (2, 88) = 0.5557 | 0.5757 | ns | |
| | | | | | | | Time x Drug | F (2, 88) = 1.048 | 0.3550 | ns | |
| | | | | | | | Treatment x Drug | F (1, 44) = 0.07768 | 0.7818 | ns | |
| | | | | | | | Time x Treatment x Drug | F (2, 88) = 0.3509 | 0.7050 | ns | |
| | | | | | 2way RMANOVA | Vehicle | Time x Drug | F (2, 40) = 0.1320 | 0.8767 | ns | 3L |
| | | | | | | | **Time** | **F (1.825, 36.49) = 9.718** | **0.0006** | ******* | |
| | | | | | | | Drug | F (1, 20) = 0.0003502 | 0.9853 | ns | |
| | | | | | | Tamoxifen | Time x Drug | F (2, 48) = 1.814 | 0.1740 | ns | 3M |
| | | | | | | | **Time** | **F (1.721, 41.30) = 12.23** | **0.0001** | ******* | |
| | | | | | | | Drug | F (1, 24) = 0.1129 | 0.7398 | ns | |
| | | | | Average freezing (%) | 2way ANOVA | - | Treatment x Drug | F (1, 44) = 0.07768 | 0.7818 | ns | 3N |
| | | | | | | | Treatment | F (1, 44) = 0.2584 | 0.6138 | ns | |
| | | | | | | | Drug | F (1, 44) = 0.06718 | 0.7967 | ns | |
| | | | CFC Day 3 | Freezing per min (%) | 3way RMANOVA | - | **Time** | **F (1.762, 77.51) = 20.89** | **<0.0001** | ******** | 3O-P |
| | | | | | | | Treatment | F (1, 44) = 0.3910 | 0.5350 | ns | |
| | | | | | | | Drug | F (1, 44) = 2.679 | 0.1088 | ns | |
| | | | | | | | Time x Treatment | F (2, 88) = 0.1470 | 0.8635 | ns | |
| | | | | | | | Time x Drug | F (2, 88) = 1.489 | 0.2313 | ns | |
| | | | | | | | Treatment x Drug | F (1, 44) = 1.934 | 0.1714 | ns | |
| | | | | | | | Time x Treatment x Drug | F (2, 88) = 0.4832 | 0.6184 | ns | |
| | | | | | 2way RMANOVA | Vehicle | Time x Drug | F (2, 40) = 1.325 | 0.2772 | ns | 3O |
| | | | | | | | **Time** | **F (1.602, 32.03) = 6.865** | **0.0056** | ****** | |
| | | | | | | | **Drug** | **F (1, 20) = 6.482** | **0.0192** | ***** | |
| | | | | | | Tamoxifen | Time x Drug | F (2, 48) = 0.3260 | 0.7234 | ns | 3P |
| | | | | | | | **Time** | **F (1.640, 39.37) = 15.86** | **<0.0001** | ******** | |
| | | | | | | | Drug | F (1, 24) = 0.02593 | 0.8734 | ns | |
| | | | | Average freezing (%) | 2way ANOVA | - | Treatment x Drug | F (1, 44) = 1.934 | 0.1714 | ns | 3Q |
| | | | | | | | Treatment | F (1, 44) = 0.3910 | 0.5350 | ns | |
| | | | | | | | Drug | F (1, 44) = 2.679 | 0.1088 | ns | |

| Genetic Line / Cohort | Strain | Sex | Behavioral Paradigm | Measurement | Statistical Test | Group | Comparison | F, t, or χ2 | p | * | Fig. |
|---|---|---|---|---|---|---|---|---|---|---|---|

| | | | | | | | | | | | |
|---|---|---|---|---|---|---|---|---|---|---|---|
| GFAP-TK | C57BL/6J | M | FST Day 1 | Immobility time per min (sec) | 3way RMANOVA | - | **Time** | **F (5, 80) = 8.951** | **<0.0001** | **** | S4A-B |
| | | | | | | | Genotype | F (1, 16) = 0.1839 | 0.6738 | ns | |
| | | | | | | | Drug | F (1, 16) = 1.959 | 0.1807 | ns | |
| | | | | | | | Time x Genotype | F (5, 80) = 0.7353 | 0.5992 | ns | |
| | | | | | | | Time x Drug | F (5, 80) = 1.051 | 0.3938 | ns | |
| | | | | | | | Genotype x Drug | F (1, 16) = 0.02947 | 0.8659 | ns | |
| | | | | | | | Time x Genotype x Drug | F (5, 80) = 0.8650 | 0.5084 | ns | |
| | | | | | 2way RMANOVA | Wild-type | Time x Drug | F (5, 35) = 1.217 | 0.3217 | ns | S4A |
| | | | | | | | **Time** | **F (2.448, 17.13) = 4.607** | **0.0199** | * | |
| | | | | | | | Drug | F (1, 7) = 1.340 | 0.2850 | ns | |
| | | | | | | GFAP-TK | Time x Drug | F (5, 45) = 0.6850 | 0.6372 | ns | S4B |
| | | | | | | | **Time** | **F (2.661, 23.95) = 5.208** | **0.0082** | ** | |
| | | | | | | | Drug | F (1, 9) = 0.7447 | 0.4106 | ns | |
| | | | | Average immobility time (sec) | 2way ANOVA | - | Genotype x Drug | F (1, 16) = 0.01959 | 0.8904 | ns | S4C |
| | | | | | | | Genotype | F (1, 16) = 0.09401 | 0.7631 | ns | |
| | | | | | | | Drug | F (1, 16) = 2.311 | 0.1480 | ns | |
| | | | FST Day 2 | Immobility time per min (sec) | 3way RMANOVA | - | Time | F (3.196, 70.30) = 2.107 | 0.1032 | ns | 4D-E |
| | | | | | | | Genotype | F (1, 22) = 1.135 | 0.2983 | ns | |
| | | | | | | | Drug | F (1, 22) = 0.3871 | 0.5402 | ns | |
| | | | | | | | Time x Genotype | F (5, 110) = 1.907 | 0.0989 | ns | |
| | | | | | | | Time x Drug | F (5, 110) = 0.3933 | 0.8525 | ns | |
| | | | | | | | Genotype x Drug | F (1, 22) = 1.154 | 0.2943 | ns | |
| | | | | | | | Time x Genotype x Drug | F (5, 110) = 0.5352 | 0.7492 | ns | |
| | | | | | 2way RMANOVA | Wild-type | Time x Drug | F (5, 45) = 0.8968 | 0.4915 | ns | 4D |
| | | | | | | | Time | F (3.287, 29.59) = 1.357 | 0.2745 | ns | |
| | | | | | | | Drug | F (1, 9) = 0.06777 | 0.8005 | ns | |
| | | | | | | GFAP-TK | Time x Drug | F (5, 65) = 0.2688 | 0.9285 | ns | 4E |
| | | | | | | | Time | F (2.833, 36.83) = 2.807 | 0.0558 | ns | |
| | | | | | | | Drug | F (1, 13) = 2.156 | 0.1658 | ns | |
| | | | | Average immobility time (sec) | 2way ANOVA | - | Genotype x Drug | F (1, 22) = 1.223 | 0.2808 | ns | 4F |
| | | | | | | | Genotype | F (1, 22) = 1.300 | 0.2665 | ns | |
| | | | | | | | Drug | F (1, 22) = 0.4900 | 0.4913 | ns | |
| | | | EPM | Time in open arms (sec) | 2way ANOVA | - | Genotype x Drug | F (1, 24) = 2.728 | 0.1117 | ns | 4G |
| | | | | | | | Genotype | F (1, 24) = 0.09373 | 0.7621 | ns | |
| | | | | | | | Drug | F (1, 24) = 3.958 | 0.0582 | ns | |
| | | | | Time in closed arms (sec) | 2way ANOVA | - | Genotype x Drug | F (1, 24) = 2.145 | 0.1560 | ns | S4D |
| | | | | | | | Genotype | F (1, 24) = 0.1592 | 0.6935 | ns | |
| | | | | | | | Drug | F (1, 24) = 3.725 | 0.0655 | ns | |
| | | | NSF | Fraction of mice not eating | Kaplan-Meier | - | Genotype x Drug | $\chi^2$ = 4.153 | 0.2454 | ns | 4H-I |
| | | | | | | Wild-type | Drug | $\chi^2$ = 0.02604 | 0.8718 | ns | 4H |
| | | | | | | GFAP-TK | Drug | $\chi^2$ = 0.7939 | 0.3729 | ns | 4I |
| | | | | Latency to feed in arena (sec) | 2way ANOVA | - | Genotype x Drug | F (1, 26) = 0.1917 | 0.6651 | ns | 4J |
| | | | | | | | Genotype | F (1, 26) = 3.105 | 0.0898 | ns | |
| | | | | | | | Drug | F (1, 26) = 0.8541 | 0.3639 | ns | |
| | | | | Weight lost (g) | 2way ANOVA | - | Genotype x Drug | F (1, 26) = 0.8560 | 0.3634 | ns | 4K |
| | | | | | | | Genotype | F (1, 26) = 1.712 | 0.2021 | ns | |
| | | | | | | | Drug | F (1, 26) = 2.591 | 0.1195 | ns | |
| | | | CFC Day 1 | Freezing per min (%) | 3way RMANOVA | - | Time | F (1.687, 43.85) = 2.866 | 0.0760 | ns | S4E-F |
| | | | | | | | **Genotype** | **F (1, 26) = 7.389** | **0.0115** | * | |
| | | | | | | | Drug | F (1, 26) = 0.06130 | 0.8064 | ns | |
| | | | | | | | Time x Genotype | F (2, 52) = 0.4733 | 0.6256 | ns | |
| | | | | | | | Time x Drug | F (2, 52) = 0.9284 | 0.4016 | ns | |
| | | | | | | | Genotype x Drug | F (1, 26) = 1.752 | 0.1972 | ns | |
| | | | | | | | Time x Genotype x Drug | F (2, 52) = 0.7129 | 0.4950 | ns | |
| | | | | | 2way RMANOVA | Wild-type | Time x Drug | F (2, 22) = 2.510 | 0.1042 | ns | S4E |
| | | | | | | | Time | F (1.179, 12.97) = 2.332 | 0.1489 | ns | |
| | | | | | | | Drug | F (1, 11) = 1.816 | 0.2049 | ns | |
| | | | | | | GFAP-TK | Time x Drug | F (2, 30) = 0.6983 | 0.5053 | ns | S4F |
| | | | | | | | Time | F (1.692, 25.38) = 1.999 | 0.1614 | ns | |
| | | | | | | | Drug | F (1, 15) = 0.9336 | 0.3492 | ns | |
| | | | | Average freezing (%) | 2way ANOVA | - | Genotype x Drug | F (1, 28) = 1.787 | 0.1920 | ns | S4G |
| | | | | | | | **Genotype** | **F (1, 28) = 7.123** | **0.0125** | * | |
| | | | | | | | Drug | F (1, 28) = 0.09585 | 0.7592 | ns | |
| | | | | | Šidák's | Saline | Genotype | t = 0.9117 | 0.6027 | ns | |
| | | | | | | (R,S)-ketamine | **Genotype** | **t = 2.933** | **0.0132** | * | |
| | | | CFC Day 2 | Freezing per min (%) | 3way RMANOVA | - | Time | F (1.724, 44.82) = 2.543 | 0.0972 | ns | 4L-M |
| | | | | | | | Genotype | F (1, 26) = 1.369 | 0.2527 | ns | |
| | | | | | | | Drug | F (1, 26) = 0.01485 | 0.9040 | ns | |
| | | | | | | | Time x Genotype | F (2, 52) = 0.5346 | 0.5891 | ns | |
| | | | | | | | Time x Drug | F (2, 52) = 0.3652 | 0.6958 | ns | |
| | | | | | | | Genotype x Drug | F (1, 26) = 1.020 | 0.3219 | ns | |
| | | | | | | | Time x Genotype x Drug | F (2, 52) = 0.5046 | 0.6067 | ns | |
| | | | | | 2way RMANOVA | Wild-type | Time x Drug | F (2, 22) = 0.8156 | 0.4553 | ns | 4L |
| | | | | | | | Time | F (1.521, 16.73) = 1.371 | 0.2742 | ns | |
| | | | | | | | Drug | F (1, 11) = 0.3844 | 0.5479 | ns | |
| | | | | | | GFAP-TK | Time x Drug | F (2, 30) = 0.3452 | 0.7109 | ns | 4M |
| | | | | | | | Time | F (1.622, 24.33) = 1.945 | 0.1702 | ns | |
| | | | | | | | Drug | F (1, 15) = 0.6938 | 0.4179 | ns | |
| | | | | Average freezing (%) | 2way ANOVA | - | Genotype x Drug | F (1, 26) = 1.020 | 0.3219 | ns | 4N |
| | | | | | | | Genotype | F (1, 26) = 1.369 | 0.2527 | ns | |
| | | | | | | | Drug | F (1, 26) = 0.01485 | 0.9040 | ns | |
| | | | CFC Day 3 | Freezing per min (%) | 3way RMANOVA | - | **Time** | **F (1.946, 50.60) = 13.56** | **<0.0001** | **** | 4O-P |
| | | | | | | | **Genotype** | **F (1, 26) = 7.754** | **0.0099** | ** | |
| | | | | | | | Drug | F (1, 26) = 2.565 | 0.1214 | ns | |
| | | | | | | | Time x Genotype | F (2, 52) = 2.017 | 0.1433 | ns | |
| | | | | | | | Time x Drug | F (2, 52) = 1.059 | 0.3541 | ns | |
| | | | | | | | Genotype x Drug | F (1, 26) = 0.6996 | 0.4105 | ns | |
| | | | | | | | Time x Genotype x Drug | F (2, 52) = 0.1413 | 0.8686 | ns | |
| | | | | | 2way RMANOVA | Wild-type | Time x Drug | F (2, 22) = 0.8775 | 0.4299 | ns | 4O |
| | | | | | | | **Time** | **F (1.431, 15.74) = 10.71** | **0.0024** | ** | |
| | | | | | | | Drug | F (1, 11) = 0.1397 | 0.7157 | ns | |
| | | | | | | GFAP-TK | Time x Drug | F (2, 30) = 0.3729 | 0.6919 | ns | 4P |
| | | | | | | | **Time** | **F (1.581, 23.72) = 5.509** | **0.0156** | * | |
| | | | | | | | **Drug** | **F (1, 15) = 9.061** | **0.0088** | ** | |
| | | | | Average freezing (%) | 2way ANOVA | - | Genotype x Drug | F (1, 26) = 0.6996 | 0.4105 | ns | 4Q |
| | | | | | | | **Genotype** | **F (1, 26) = 7.754** | **0.0099** | ** | |
| | | | | | | | Drug | F (1, 26) = 2.565 | 0.1214 | ns | |
| | | | | | Šidák's | Saline | **Genotype** | **t = 2.537** | **0.0175** | * | |
| | | | | | | (R,S)-ketamine | Genotype | t = 1.39 | 0.1762 | ns | |
| | | | | | unpaired t-test | GFAP-TK | **Drug** | **t = 3.010** | **0.0088** | ** | |

| Genetic Line / Cohort | Strain | Sex | Behavioral Paradigm | Measurement | Statistical Test | Group | Comparison | F, t, or $\chi^2$ | p | * | Fig. |
|---|---|---|---|---|---|---|---|---|---|---|---|
| | | | | | | | **Time** | **F (5, 180) = 7.123** | **<0.0001** | **** | |
| | | | | | | | Genotype | F (1, 36) = 0.2002 | 0.6572 | ns | |
| | | | | | | | **Drug** | **F (1, 36) = 16.13** | **0.0003** | *** | |

| | | | | | | | | | | | |
|---|---|---|---|---|---|---|---|---|---|---|---|
| PVCre | 129S6/SvEv | M | FST Day 1 | Immobility time per min (sec) | 3way RMANOVA | - | Time x Genotype | F (5, 180) = 0.7560 | 0.5827 | ns | S5A-B |
| | | | | | | | Time x Drug | F (5, 180) = 0.8357 | 0.5259 | ns | |
| | | | | | | | Genotype x Drug | F (1, 36) = 4.261 | 0.0463 | * | |
| | | | | | | | Time x Genotype x Drug | F (5, 180) = 0.6624 | 0.6524 | ns | |
| | | | | | 2way RMANOVA | PVCre(-) | Time x Drug | F (5, 75) = 0.9416 | 0.4593 | ns | S5A |
| | | | | | | | Time | F (3.389, 50.83) = 4.874 | 0.0034 | ** | |
| | | | | | | | Drug | F (1, 15) = 1.152 | 0.3001 | ns | |
| | | | | | | PVCre(+) | Time x Drug | F (5, 105) = 0.5704 | 0.7225 | ns | S5B |
| | | | | | | | Time | F (3.318, 69.68) = 3.090 | 0.0283 | * | |
| | | | | | | | Drug | F (1, 21) = 31.68 | <0.0001 | **** | |
| | | | | Average immobility time (sec) | 2way ANOVA | - | Genotype x Drug | F (1, 36) = 3.505 | 0.0693 | ns | S5C |
| | | | | | | | Genotype | F (1, 36) = 0.01202 | 0.9133 | ns | |
| | | | | | | | Drug | F (1, 36) = 15.84 | 0.0003 | *** | |
| | | | | | Šidák's | PVCre(-) | Drug | t = 1.389 | 0.3167 | ns | |
| | | | | | | PVCre(+) | Drug | t = 4.49 | 0.0001 | *** | |
| | | | FST Day 2 | Immobility time per min (sec) | 3way RMANOVA | - | Time | F (3.208, 112.3) = 0.7386 | 0.5396 | ns | 5D-E |
| | | | | | | | Genotype | F (1, 35) = 0.3370 | 0.5653 | ns | |
| | | | | | | | Drug | F (1, 35) = 6.978 | 0.0123 | * | |
| | | | | | | | Time x Genotype | F (5, 175) = 0.6707 | 0.6462 | ns | |
| | | | | | | | Time x Drug | F (5, 175) = 0.2400 | 0.9443 | ns | |
| | | | | | | | Genotype x Drug | F (1, 35) = 1.207 | 0.2794 | ns | |
| | | | | | | | Time x Genotype x Drug | F (5, 175) = 0.3561 | 0.8778 | ns | |
| | | | | | 2way RMANOVA | PVCre(-) | Time x Drug | F (5, 75) = 0.1460 | 0.9806 | ns | 5D |
| | | | | | | | Time | F (2.976, 44.64) = 1.017 | 0.3939 | ns | |
| | | | | | | | Drug | F (1, 15) = 0.8880 | 0.3610 | ns | |
| | | | | | | PVCre(+) | Time x Drug | F (5, 100) = 0.4951 | 0.7793 | ns | 5E |
| | | | | | | | Time | F (2.902, 58.04) = 0.3016 | 0.8177 | ns | |
| | | | | | | | Drug | F (1, 20) = 9.345 | 0.0062 | ** | |
| | | | | Average immobility time (sec) | 2way ANOVA | - | Genotype x Drug | F (1, 35) = 1.246 | 0.2719 | ns | 5F |
| | | | | | | | Genotype | F (1, 35) = 0.5691 | 0.4557 | ns | |
| | | | | | | | Drug | F (1, 35) = 5.472 | 0.0252 | * | |
| | | | | | Šidák's | PVCre(-) | Drug | t = 0.8135 | 0.6653 | ns | |
| | | | | | | PVCre(+) | Drug | t = 2.619 | 0.0257 | * | |
| | | | EPM | Time in open arms (sec) | 2way ANOVA | - | Genotype x Drug | F (1, 36) = 0.02196 | 0.8830 | ns | 5G |
| | | | | | | | Genotype | F (1, 36) = 3.224 | 0.0810 | ns | |
| | | | | | | | Drug | F (1, 36) = 0.1271 | 0.7235 | ns | |
| | | | | Time in closed arms (sec) | 2way ANOVA | - | Genotype x Drug | F (1, 36) = 0.8546 | 0.3614 | ns | S5D |
| | | | | | | | Genotype | F (1, 36) = 5.774 | 0.0215 | * | |
| | | | | | | | Drug | F (1, 36) = 1.614 | 0.2120 | ns | |
| | | | | | Šidák's | Saline | Genotype | t = 2.417 | 0.0413 | * | |
| | | | | | | (R,S)-ketamine | Genotype | t = 1.019 | 0.5307 | ns | |
| | | | NSF | Fraction of mice not eating | Kaplan-Meier | - | Genotype x Drug | $\chi^2$ = 1.03 | 0.7941 | ns | 5H-I |
| | | | | | | PVCre(-) | Drug | $\chi^2$ = 0.5925 | 0.4415 | ns | 5H |
| | | | | | | PVCre(+) | Drug | $\chi^2$ = 0.2238 | 0.6362 | ns | 5I |
| | | | | Latency to feed in arena (sec) | 2way ANOVA | - | Genotype x Drug | F (1, 35) = 0.002354 | 0.9616 | ns | 5J |
| | | | | | | | Genotype | F (1, 35) = 0.7042 | 0.4071 | ns | |
| | | | | | | | Drug | F (1, 35) = 0.4527 | 0.5055 | ns | |
| | | | | Weight lost (g) | 2way ANOVA | - | Genotype x Drug | F (1, 35) = 0.9286 | 0.3418 | ns | 5K |
| | | | | | | | Genotype | F (1, 35) = 0.4064 | 0.5280 | ns | |
| | | | | | | | Drug | F (1, 35) = 4.074 | 0.0513 | ns | |
| | | | CFC Day 1 | Freezing per min (%) | 3way RMANOVA | - | Time | F (1.865, 67.15) = 7.146 | 0.0019 | ** | S5E-F |
| | | | | | | | Genotype | F (1, 36) = 1.433 | 0.2392 | ns | |
| | | | | | | | Drug | F (1, 36) = 1.146 | 0.2914 | ns | |
| | | | | | | | Time x Genotype | F (2, 72) = 2.142 | 0.1248 | ns | |
| | | | | | | | Time x Drug | F (2, 72) = 2.532 | 0.0866 | ns | |
| | | | | | | | Genotype x Drug | F (1, 36) = 0.2297 | 0.6346 | ns | |
| | | | | | | | Time x Genotype x Drug | F (2, 72) = 0.8689 | 0.4238 | ns | |
| | | | | | 2way RMANOVA | PVCre(-) | Time x Drug | F (2, 30) = 2.578 | 0.0926 | ns | S5E |
| | | | | | | | Time | F (1.985, 29.77) = 0.6927 | 0.5070 | ns | |
| | | | | | | | Drug | F (1, 15) = 0.2730 | 0.6090 | ns | |
| | | | | | | PVCre(+) | Time x Drug | F (2, 42) = 0.6327 | 0.5361 | ns | S5F |
| | | | | | | | Time | F (1.642, 34.49) = 9.757 | 0.0009 | *** | |
| | | | | | | | Drug | F (1, 21) = 1.074 | 0.3118 | ns | |
| | | | | Average freezing (%) | 2way ANOVA | - | Genotype x Drug | F (1, 36) = 0.2297 | 0.6347 | ns | S5G |
| | | | | | | | Genotype | F (1, 36) = 1.432 | 0.2392 | ns | |
| | | | | | | | Drug | F (1, 36) = 1.146 | 0.2916 | ns | |
| | | | CFC Day 2 | Freezing per min (%) | 3way RMANOVA | - | Time | F (1.795, 64.63) = 32.68 | <0.0001 | **** | 5L-M |
| | | | | | | | Genotype | F (1, 36) = 0.1043 | 0.7486 | ns | |
| | | | | | | | Drug | F (1, 36) = 2.962 | 0.0938 | ns | |
| | | | | | | | Time x Genotype | F (2, 72) = 0.3863 | 0.6810 | ns | |
| | | | | | | | Time x Drug | F (2, 72) = 4.449 | 0.0151 | * | |
| | | | | | | | Genotype x Drug | F (1, 36) = 3.445 | 0.0716 | ns | |
| | | | | | | | Time x Genotype x Drug | F (2, 72) = 0.2531 | 0.7771 | ns | |
| | | | | | 2way RMANOVA | PVCre(-) | Time x Drug | F (2, 30) = 2.927 | 0.0690 | ns | 5L |
| | | | | | | | Time | F (1.692, 25.38) = 17.38 | <0.0001 | **** | |
| | | | | | | | Drug | F (1, 15) = 6.094 | 0.0261 | * | |
| | | | | | | PVCre(+) | Time x Drug | F (2, 42) = 1.612 | 0.2117 | ns | 5M |
| | | | | | | | Time | F (1.657, 34.80) = 15.58 | <0.0001 | **** | |
| | | | | | | | Drug | F (1, 21) = 0.01009 | 0.9209 | ns | |
| | | | | Average freezing (%) | 2way ANOVA | - | Genotype x Drug | F (1, 36) = 3.445 | 0.0716 | ns | 5N |
| | | | | | | | Genotype | F (1, 36) = 0.1043 | 0.7486 | ns | |
| | | | | | | | Drug | F (1, 36) = 2.962 | 0.0938 | ns | |
| | | | CFC Day 3 | Freezing per min (%) | 3way RMANOVA | - | Time | F (1.869, 67.27) = 55.28 | <0.0001 | **** | 5O-P |
| | | | | | | | Genotype | F (1, 36) = 0.003695 | 0.9519 | ns | |
| | | | | | | | Drug | F (1, 36) = 3.056 | 0.0890 | ns | |
| | | | | | | | Time x Genotype | F (2, 72) = 0.2522 | 0.7778 | ns | |
| | | | | | | | Time x Drug | F (2, 72) = 1.325 | 0.2723 | ns | |
| | | | | | | | Genotype x Drug | F (1, 36) = 1.535e-005 | 0.9969 | ns | |
| | | | | | | | Time x Genotype x Drug | F (2, 72) = 1.450 | 0.2414 | ns | |
| | | | | | 2way RMANOVA | PVCre(-) | Time x Drug | F (2, 30) = 2.286 | 0.1191 | ns | 5O |
| | | | | | | | Time | F (1.872, 28.08) = 25.07 | <0.0001 | **** | |
| | | | | | | | Drug | F (1, 15) = 1.126 | 0.3054 | ns | |
| | | | | | | PVCre(+) | Time x Drug | F (2, 42) = 0.03021 | 0.9703 | ns | 5P |
| | | | | | | | Time | F (1.727, 36.28) = 30.88 | <0.0001 | **** | |
| | | | | | | | Drug | F (1, 21) = 2.064 | 0.1655 | ns | |
| | | | | Average freezing (%) | 2way ANOVA | - | Genotype x Drug | F (1, 36) = 1.531e-005 | 0.9969 | ns | 5Q |
| | | | | | | | Genotype | F (1, 36) = 0.003695 | 0.9519 | ns | |
| | | | | | | | Drug | F (1, 36) = 3.056 | 0.0890 | ns | |

| Genetic Line / Cohort | Strain | Sex | Behavioral Paradigm | Measurement | Statistical Test | Group | Comparison | F, t, or $\chi^2$ | p | * | Fig. |
|---|---|---|---|---|---|---|---|---|---|---|---|
| | | | | | 3way RMANOVA (mixed-effects anaysis) | - | Time | F (5, 213) = 2.409 | 0.0377 | * | S7A-B |
| | | | | | | | Genotype | F (1, 45) = 5.557 | 0.0228 | * | |
| | | | | | | | Drug | F (1, 45) = 9.331 | 0.0038 | ** | |
| | | | | | | | Time x Genotype | F (5, 213) = 1.046 | 0.3919 | ns | |
| | | | | | | | Time x Drug | F (5, 213) = 0.4420 | 0.8188 | ns | |

| | | | | | | | | | | | |
|---|---|---|---|---|---|---|---|---|---|---|---|
| PVCre | 129S6/SvEv | F | FST Day 1 | Immobility time per min (sec) | | | Genotype x Drug | F (1, 45) = 0.2508 | 0.6189 | ns | |
| | | | | | | | Time x Genotype x Drug | F (5, 213) = 0.1854 | 0.9679 | ns | |
| | | | | | 2way RMANOVA (mixed-effects analysis) | PVCre(-) | Time x Drug | F (5, 104) = 0.3562 | 0.8772 | ns | S7A |
| | | | | | | | Time | F (3.801, 79.05) = 0.8108 | 0.5166 | ns | |
| | | | | | | | Drug | F (1, 22) = 3.801 | 0.0641 | ns | |
| | | | | | | PVCre(+) | Time x Drug | F (5, 109) = 0.2855 | 0.9202 | ns | S7B |
| | | | | | | | Time | F (2.855, 62.25) = 2.525 | 0.0685 | ns | |
| | | | | | | | **Drug** | **F (1, 23) = 5.723** | **0.0253** | ***** | |
| | | | | Average immobility time (sec) | 2way ANOVA | - | Genotype x Drug | F (1, 45) = 0.3687 | 0.5468 | ns | S7C |
| | | | | | | | **Treatment** | **F (1, 45) = 5.795** | **0.0202** | ***** | |
| | | | | | | | **Drug** | **F (1, 45) = 8.752** | **0.0049** | ****** | |
| | | | | | Šidák's | PVCre(-) | Drug | t = 1.601 | 0.2192 | ns | |
| | | | | | | PVCre(+) | **Drug** | **t = 2.626** | **0.0234** | ***** | |
| | | | | | Šidák's | Saline | Treatment | t = 1.947 | 0.1122 | ns | |
| | | | | | | (R,S)-ketamine | Treatment | t = 1.421 | 0.2979 | ns | |
| | | | FST Day 2 | Immobility time per min (sec) | 3way RMANOVA (mixed-effects anaysis) | - | Time | F (4.040, 162.4) = 1.425 | 0.2275 | ns | S6D-E |
| | | | | | | | Genotype | F (1, 45) = 3.770 | 0.0585 | ns | |
| | | | | | | | Drug | F (1, 45) = 0.8194 | 0.3702 | ns | |
| | | | | | | | Time x Genotype | F (5, 201) = 0.7768 | 0.5674 | ns | |
| | | | | | | | Time x Drug | F (5, 201) = 0.5030 | 0.7738 | ns | |
| | | | | | | | Genotype x Drug | F (1, 45) = 0.1164 | 0.7345 | ns | |
| | | | | | | | Time x Genotype x Drug | F (5, 201) = 0.5665 | 0.7257 | ns | |
| | | | | | 2way RMANOVA (mixed-effects analysis) | PVCre(-) | Time x Drug | F (5, 92) = 0.4776 | 0.7921 | ns | S6D |
| | | | | | | | Time | F (3.608, 66.39) = 1.485 | 0.2206 | ns | |
| | | | | | | | Drug | F (1, 22) = 0.1284 | 0.7236 | ns | |
| | | | | | | PVCre(+) | Time x Drug | F (5, 109) = 0.5936 | 0.7049 | ns | S6E |
| | | | | | | | Time | F (3.211, 70.00) = 0.5133 | 0.6867 | ns | |
| | | | | | | | Drug | F (1, 23) = 0.9769 | 0.3332 | ns | |
| | | | | Average immobility time (sec) | 2way ANOVA | - | Genotype x Drug | F (1, 45) = 0.001652 | 0.9678 | ns | S6F |
| | | | | | | | Treatment | F (1, 45) = 3.686 | 0.0612 | ns | |
| | | | | | | | Drug | F (1, 45) = 0.9104 | 0.3451 | ns | |
| | | | EPM | Time in open arms (sec) | 2way ANOVA | - | Genotype x Drug | F (1, 45) = 0.3351 | 0.5656 | ns | S6G |
| | | | | | | | Treatment | F (1, 45) = 0.2578 | 0.6141 | ns | |
| | | | | | | | Drug | F (1, 45) = 0.08917 | 0.7666 | ns | |
| | | | | Time in closed arms (sec) | 2way ANOVA | - | Genotype x Drug | F (1, 45) = 0.2507 | 0.6190 | ns | S7D |
| | | | | | | | Treatment | F (1, 45) = 0.3283 | 0.5695 | ns | |
| | | | | | | | Drug | F (1, 45) = 0.1604 | 0.6906 | ns | |
| | | | NSF | Fraction of mice not eating | Kaplan-Meier | - | Genotype x Drug | $\chi2$ = 6.496 | 0.0898 | ns | S6H-I |
| | | | | | | PVCre(-) | **Drug** | **$\chi2$ = 4.65** | **0.0311** | ***** | S6H |
| | | | | | | PVCre(+) | Drug | $\chi2$ = 1.052 | 0.3051 | ns | S6I |
| | | | | Latency to feed in arena (sec) | 2way ANOVA | - | Genotype x Drug | F (1, 45) = 1.675 | 0.2022 | ns | S6J |
| | | | | | | | Treatment | F (1, 45) = 2.051 | 0.1590 | ns | |
| | | | | | | | **Drug** | **F (1, 45) = 5.551** | **0.0229** | ***** | |
| | | | | | Šidák's | PVCre(-) | **Drug** | **t = 2.486** | **0.0332** | ***** | |
| | | | | | | PVCre(+) | Drug | t = 0.7822 | 0.6844 | ns | |
| | | | | Weight lost (g) | 2way ANOVA | - | Genotype x Drug | F (1, 45) = 0.001659 | 0.9677 | ns | S6K |
| | | | | | | | Treatment | F (1, 45) = 0.2244 | 0.6380 | ns | |
| | | | | | | | Drug | F (1, 45) = 0.01729 | 0.8960 | ns | |
| | | | CFC Day 1 | Freezing per min (%) | 3way RMANOVA | - | **Time** | **F (1.735, 74.61) = 3.930** | **0.0291** | ***** | S7E-F |
| | | | | | | | Genotype | F (1, 43) = 1.132 | 0.2934 | ns | |
| | | | | | | | Drug | F (1, 43) = 0.05941 | 0.8086 | ns | |
| | | | | | | | Time x Genotype | F (2, 86) = 0.4566 | 0.6350 | ns | |
| | | | | | | | **Time x Drug** | **F (2, 86) = 6.090** | **0.0034** | ****** | |
| | | | | | | | Genotype x Drug | F (1, 43) = 2.384 | 0.1299 | ns | |
| | | | | | | | Time x Genotype x Drug | F (2, 86) = 0.2880 | 0.7505 | ns | |
| | | | | | 2way RMANOVA | PVCre(-) | **Time x Drug** | **F (2, 44) = 4.275** | **0.0201** | ***** | S7E |
| | | | | | | | Time | F (1.507, 33.15) = 2.748 | 0.0918 | ns | |
| | | | | | | | Drug | F (1, 22) = 0.8171 | 0.3758 | ns | |
| | | | | | Šidák's | PVCre(-) | Minute 1 Drug | t = 1.846 | 0.2795 | ns | |
| | | | | | | | Minute 2 Drug | t = 1.251 | 0.5699 | ns | |
| | | | | | | | Minute 3 Drug | t = 1.497 | 0.3832 | ns | |
| | | | | | 2way RMANOVA | PVCre(+) | Time x Drug | F (2, 42) = 2.494 | 0.0947 | ns | S7F |
| | | | | | | | Time | F (1.856, 38.98) = 1.840 | 0.1746 | ns | |
| | | | | | | | Drug | F (1, 21) = 1.655 | 0.2123 | ns | |
| | | | | Average freezing (%) | 2way ANOVA | - | Genotype x Drug | F (1, 43) = 2.385 | 0.1298 | ns | S7G |
| | | | | | | | Treatment | F (1, 43) = 1.132 | 0.2932 | ns | |
| | | | | | | | Drug | F (1, 43) = 0.05938 | 0.8086 | ns | |
| | | | CFC Day 2 | Freezing per min (%) | 3way RMANOVA | - | **Time** | **F (1.918, 82.48) = 36.50** | **<0.0001** | ******** | S6L-M |
| | | | | | | | Genotype | F (1, 43) = 1.389 | 0.2450 | ns | |
| | | | | | | | Drug | F (1, 43) = 2.385 | 0.1299 | ns | |
| | | | | | | | Time x Genotype | F (2, 86) = 0.9910 | 0.3754 | ns | |
| | | | | | | | Time x Drug | F (2, 86) = 0.8566 | 0.4282 | ns | |
| | | | | | | | Genotype x Drug | F (1, 43) = 2.924 | 0.0945 | ns | |
| | | | | | | | Time x Genotype x Drug | F (2, 86) = 0.3601 | 0.6987 | ns | |
| | | | | | 2way RMANOVA | PVCre(-) | Time x Drug | F (2, 44) = 0.3445 | 0.7104 | ns | S6L |
| | | | | | | | **Time** | **F (1.953, 42.97) = 15.82** | **<0.0001** | ******** | |
| | | | | | | | **Drug** | **F (1, 22) = 5.201** | **0.0326** | ***** | |
| | | | | | | PVCre(+) | Time x Drug | F (2, 42) = 0.9030 | 0.4131 | ns | S6M |
| | | | | | | | **Time** | **F (1.844, 38.73) = 22.00** | **<0.0001** | ******** | |
| | | | | | | | Drug | F (1, 21) = 0.01402 | 0.9069 | ns | |
| | | | | Average freezing (%) | 2way ANOVA | - | Genotype x Drug | F (1, 43) = 2.925 | 0.0944 | ns | S6N |
| | | | | | | | Treatment | F (1, 43) = 1.389 | 0.2450 | ns | |
| | | | | | | | Drug | F (1, 43) = 2.385 | 0.1299 | ns | |
| | | | CFC Day 3 | Freezing per min (%) | 3way RMANOVA | - | **Time** | **F (1.672, 71.91) = 38.34** | **<0.0001** | ******** | S6O-P |
| | | | | | | | Genotype | F (1, 43) = 1.061 | 0.3088 | ns | |
| | | | | | | | Drug | F (1, 43) = 1.334 | 0.2544 | ns | |
| | | | | | | | Time x Genotype | F (2, 86) = 0.09550 | 0.9090 | ns | |
| | | | | | | | Time x Drug | F (2, 86) = 1.483 | 0.2328 | ns | |
| | | | | | | | Genotype x Drug | F (1, 43) = 1.315 | 0.2578 | ns | |
| | | | | | | | Time x Genotype x Drug | F (2, 86) = 2.162 | 0.1213 | ns | |
| | | | | | 2way RMANOVA | PVCre(-) | Time x Drug | F (2, 44) = 0.05165 | 0.9497 | ns | S6O |
| | | | | | | | **Time** | **F (1.746, 38.42) = 17.89** | **<0.0001** | ******** | |
| | | | | | | | Drug | F (1, 22) = 2.175 | 0.1544 | ns | |
| | | | | | | PVCre(+) | **Time x Drug** | **F (2, 42) = 3.483** | **0.0398** | ***** | S6P |
| | | | | | | | **Time** | **F (1.589, 33.37) = 20.48** | **<0.0001** | ******** | |
| | | | | | | | Drug | F (1, 21) = 4.471e-005 | 0.9947 | ns | |
| | | | | | Šidák's | PVCre(+) | Minute 1 Drug | t = 0.7637 | 0.8379 | ns | |
| | | | | | | | Minute 2 Drug | t = 0.3 | 0.9874 | ns | |
| | | | | | | | Minute 3 Drug | t = 1.111 | 0.6366 | ns | |
| | | | | Average freezing (%) | 2way ANOVA | - | Genotype x Drug | F (1, 43) = 1.315 | 0.2578 | ns | S6Q |
| | | | | | | | Treatment | F (1, 43) = 1.061 | 0.3088 | ns | |
| | | | | | | | Drug | F (1, 43) = 1.334 | 0.2544 | ns | |
