## Supplementary material for "(*R,S*)-ketamine’s rapid-acting antidepressant effects are modulated by NR2B-containing NMDA receptors on adult-born hippocampal neurons": Table 1

**Supplemental Table 2.** Key Resources Table

| Resource Type | Specific Reagent or Resources | Source or Reference | Identifiers | Additional Information |
|---|---|---|---|---|
| Antibody | Chicken polyclonal anti-GFP | Abcam, Cambridge, United Kingdom | Cat# ab13970, RRID:AB_300798 | concentration 1:500 |
| Antibody | Rabbit polyclonal anti-PV | Swant, Burgdorf, Switzerland | Cat# PV27, RRID:AB_2631173 | concentration 1:3,000 |
| Antibody | Rabbit polyclonal IgG anti-DCX | Abcam, Cambridge, United Kingdom | Cat# ab18723, RRID:AB_732011 | concentration 1:4,000 |
| Antibody | Cy2 AffiniPure donkey anti-chicken IgY (IgG) | Jackson ImunoResearch, West Grove, PA | Cat# 703-225-155, RRID:AB_2340370 | concentration 1:500 |
| Antibody | Cy3 AffiniPure donkey anti-rabbit IgG | Jackson ImunoResearch, West Grove, PA | Cat# 711-165-152, RRID:AB_2307443 | concentration 1:500 |
| Antibody | Alexa Fluor 488 AffiniPure donkey anti-rabbit IgG | Jackson ImunoResearch, West Grove, PA | Cat# 711-545-152, RRID:AB_2313584 | concentration 1:250 |
| Mounting medium | Fluoromount G | Electron Microscopy Sciences, Hatfield, PA | Cat.# 17984-25 | |
| Organism / strain | Mouse (male, female): 129S6/SvEv | Taconic Biosciences, Inc., Germantown, NY | Cat.# 129SVE; 129S6/SvEv | |
| Organism / strain | Mouse (male, female): C57BL/6J | Jackson Laboratory, Boston, MA | JAX#000664; C57BL/6J | |
| Organism / strain | Mouse (male, female): NR2B$^{f/f}$ | Jackson Laboratory, Boston, MA | JAX# 032664; C57BL/6J | doi:10.1016/j.neuron.2008.09.039. |
| Organism / strain | Mouse (male, female): EYFP$^{f/f}$ | Jackson Laboratory, Boston, MA | JAX# 006148; C57BL/6J | doi:10.1186/1471-213x-1-4. |
| Organism / strain | Mouse (male, female): NestinCreER$^{T2}$ | Jackson Laboratory, Boston, MA | JAX# 016261; C57BL/6J | doi:10.1016/j.neuron.2011.05.022. |
| Organism / strain | Mouse (male, female): PV-Cre | Jackson Laboratory, Boston, MA | JAX# 017320; C57BL/6J | doi:10.1371/journal.pbio.0030159 |
| Organism / strain | Mouse (male): GFAP-TK | Jackson Laboratory, Boston, MA | JAX# 017523; 129S6/SvEv | doi:10.1038/nature10287.; doi:10.1002/hipo.20964. |
| Compound or Drug | (*R,S*)-ketamine | Ketaset III, Fort Dodge Animal Health, Fort Dodge, IA | Cat# 0856-4403-01 | 10 or 30 mg/kg, i.p. in physiological saline |
| Compound or Drug | Tamoxifen | Sigma-Aldrich, St. Louis, MO | Cat# T5648 | 10mg/mL in 90% corn oil / 10% EtOH |
| Compound or Drug | Ganciclovir | Cytovene-IV, Roche, Indianapolis, IN | Cat# CSE-3021 | 60mg/mL in 0.9% saline |
| Compound or Drug | Hoechst 33342 | Thermo Fisher Scientific, Waltham, MA | Cat# H3570 | concentration 1:10,000 |
| Software or Algorithm | VideoTrack | ViewPoint Behavior Technology, Civrieux, France | | |

| | | |
|---|---|---|
| Software or Algorithm | ANY-maze behavior tracking | Stoelting Co., Wood Dale, IL |
| Software or Algorithm | FreezeFrame / FreezeView v4 | Actimetrics, Wilmette, IL |
| Software or Algorithm | LAS X | Leica Microsystems Inc., Wetzlar, Germany |
| Software or Algorithm | Fiji (ImageJ v.1.52p) | http://fiji.sc/; doi:10.1038/nmeth.2019 |
